# Quantifying the information about uncertainty in neural population codes

**DOI:** 10.64898/2026.07.13.738167

**Authors:** Xiaolu Wang, Peter Dayan, Paul M Bays

**Affiliations:** University of Cambridge, Department of Psychology, Cambridge, CB2 3EB, UK; Sun Yat-sen University, Department of Psychology, Guangzhou, China; Max Planck Institute for Biological Cybernetics, Tübingen, Germany; University of Tübingen, Tübingen, Germany

## Abstract

The activity of neural populations typically encodes more information about sensory or motor variables than can be captured by point estimates of the variables. We present and compare two approaches to quantifying this additional or ancillary information and its relationship to uncertainty: the mutual information between activity and estimation error, and the Fisher information loss, which can be interpreted in terms of curvature in information geometry. We show that deviations from Gaussianity of estimation errors, including the long tails frequently observed in human behavioural tasks, are an expected corollary of the presence of ancillary information. However, populations with similar distributions of estimation error can differ substantially in their ancillary information content depending on the noise characteristics. For a given population tuning and noise model, our results quantify an upper bound on the information about uncertainty that can be obtained from population activity alone: behaviour demonstrating knowledge in excess of this bound would indicate access to a separate source of information about uncertainty. Finally, we contrast the effects of external noise and decreasing internal signal strength on ancillary information and the Gaussianity of errors. Our work directly relates knowledge about uncertainty to non-Gaussianity in sensory estimates, and establishes a coherent theoretical foundation for investigating the basis of metacognition in neural population activity.

**Author summary:** The brain processes sensory evidence about the external world via inherently noisy neural activity. As a result, behavioural judgments – such as estimating the direction of a moving object – are fundamentally uncertain. While animals, including humans, routinely use uncertainty to guide decisions under risk, how neural populations represent this uncertainty remains unclear. In this work, we show how the same neural activity used to decode a sensory variable can also provide information about the estimate’s reliability. We introduce a mathematical framework to quantify this “ancillary information” directly from a neural population’s encoding model. We demonstrate that ancillary information predicts non-Gaussianity in estimation errors and sets an upper bound on metacognitive sensitivity (how accurately subjective confidence tracks performance). Crucially, we show that neural populations with distinct noise characteristics can yield near-identical estimation errors while providing very different degrees of uncertainty information. This highlights the importance of evaluating ancillary information, not just error patterns, when comparing competing models of sensory coding.

## 1 Introduction

Studies in humans and non-human primates indicate that the brain can access information about the uncertainty of internal representations to support adaptive behaviour (Bays et al., 2024; Cortese, 2022; Goris et al., 2025; Ma & Jazayeri, 2014), using it to integrate different sources of information appropriately (C.-H. S. Lin & Garrido, 2022; Murai & Yotsumoto, 2018; Wolpert & Landy, 2012) and choosing between risky actions (Bach & Dolan, 2012; Johnson & Busemeyer, 2010). For example, a driver preparing to change lanes must estimate the distance and speed of an approaching car. In daylight, when viewing conditions are good, this estimate is relatively precise, and the driver may feel the gap is sufficient and proceed. At night or in fog, the same driver may judge their estimate to be unreliable and choose to wait. In such real-world situations, decisions rely on not only an estimate of the external state, but also on information about the uncertainty of that estimate. Behavioural studies have duly shown that human subjective evaluations of uncertainty or confidence track trial-by-trial variability in estimates of continuous variables (Honig et al., 2020; Rademaker et al., 2012).

Some uncertainty is inherent in the information available to our senses (e.g. one typically cannot see what is behind one’s head), but in cases where the sensory signal is informative but weak (like seeing in fog) the uncertainty reflects internal sources of noise. A range of models have been proposed to link subjective uncertainty or confidence to statistical uncertainty in internal representations (Boundy-Singer et al., 2023; Denison et al., 2018; Li & Ma, 2020; Sanders et al., 2016). In the framework of signal detection theory (SDT), confidence in a categorical decision can be derived from the distance of a scalar internal signal from a decision boundary (Fleming & Lau, 2014; Maniscalco & Lau, 2012). However, for the continuous variables that are more frequently the focus of real-world decisions, confidence is more appropriately represented by the width or spread of the likelihood function (or the posterior distribution in a Bayesian setting) over the space of possible values.

One account of estimation for continuous variables, the variable precision (VP) model, postulates that encoding precision fluctuates randomly over objects and time because of doubly stochastic internal noise (van den Berg et al., 2012). While originally formulated to explain the presence of long tails (deviations from Gaussianity) in estimation errors in recall from visual working memory, the VP model has also been used to model confidence judgements, given the additional assumption that observers have subjective access to the encoding precision for each observation (van den Berg et al., 2017) (see also Fougnie et al. (2012)). However, the causes of precision variation are not identified by these descriptive models.

In the brain, stochasticity in neuronal spiking contributes to variability in internal representations, which gives rise to uncertainty in downstream decoding systems. However, the same activity can also carry information about uncertainty. Ma et al. (2006) showed that for Poisson spiking variability (and a broader class of “Poisson-like” noise characteristics) the summed amplitude of activity in a neural population could be used as a source of information about the uncertainty in a decoded estimate, with more activity corresponding to less uncertainty. Bays (2016) showed that patterns of error and subjective confidence in perceptual estimation of a low-contrast orientation stimulus could be simultaneously captured by a population coding model with Poisson noise characteristics. This model had previously been shown to account for long tails and the effects of memory set size and prioritization on recall errors (Bays, 2014). Schneegans et al. (2020) united the neural and descriptive accounts, showing that the VP model provides an approximation in the continuous limit to predictions of a population coding model with Poisson noise. These results suggest that, along with access to uncertainty, the long-tailed error pattern observed in continuous estimation tasks can emerge naturally from population codes.

A recent study presented an alternative, geometric perspective on such deviations from normality, describing long tails as a consequence of curvature of the encoding manifold embedded in high-dimensional neural activity space (Wei & Woodford, 2025). The underlying framework was equivalent to a population coding model with Gaussian noise (Schneegans & Bays, 2017; Schurgin et al., 2020; Tomić & Bays, 2024b), but information about uncertainty was not directly addressed in this geometric account.

Here, we present a general framework in which to situate the information about uncertainty contained in a population code, and an interpretation of it from a geometric perspective. We consider the maximum likelihood estimate (MLE) as a point estimator of a continuous variable based on activity in an idealized neural population. Information about uncertainty in an estimate is contained in *ancillary* statistics, components of the neural activity that are probabilistically independent of the estimated variable. We quantify this ancillary information using two complementary approaches: an entropy-based measure MI*e*, defined as the mutual information (MI) between neural activity and error in the MLE, and a measure based on Fisher information (FI), ℐ_loss_, calculated as the difference between FI for the neural activity versus the MLE. The latter measure is closely related to curvature in information geometry, revealing a formal relationship between encoding of uncertainty and deviations from Gaussianity in behavioural error. We investigate how the ancillary information in a population code depends on the combination of noise and tuning characteristics.

## 2 Results

### 2.1 General framework

Our main results are obtained from an encoding-decoding model where **r** is the stochastic activity of a population of neurons encoding a stimulus value *θ*, under the common assumption that the population of tuning functions has translational symmetry (we address in the Discussion how our results might be extended to cases where this symmetry does not hold). This can be understood as a specific instance of the more general problem of estimating a scalar parameter *θ* from an observation **r**, where the statistical model is of the translation-type first described by R. A. Fisher (1934), i.e. the likelihood *p*(**r** | *θ*) = *p*(*T* (**r**, *δ*) | *θ* + *δ*) ∀*θ* ∀*δ* for some transformation function *T* (**r**, ·) that is volume-preserving for **r** (see also Efron and Hinkley 1978). Estimation of a location parameter is a familiar example of this type.

The MLE 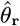 of the parameter *θ* maximizes the likelihood function over the parameter space Θ,

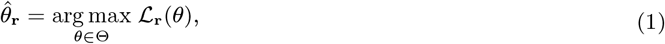

where the likelihood function ℒ_r_(*θ*) is simply *p*(**r** *θ*) treated as a function of *θ*, for a particular observation **r**. Our interest is in what an observation **r** can tell us about the reliability of its corresponding 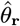 as an estimate of *θ*.

In trivial cases, 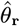 is a sufficient statistic for *θ*, in which case **r** can provide no further information about the estimator’s reliability. Where this is not the case, in a translation-type model the observation **r** can be reduced to a sufficient statistic 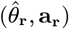 composed of the maximum likelihood estimate and a statistic **a**_**r**_ that is probabilistically independent of *θ*, but contains information about the accuracy of the MLE (R. A. Fisher, 1925). A statistic with this property of independence is described as “ancillary”, and because **a**_**r**_ is jointly sufficient with the MLE 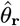 it is described as an ancillary complement.

We can decompose the likelihood as

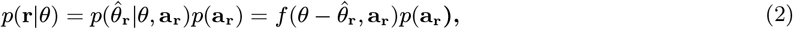

where the first step is based on the independence of **a**_**r**_ and *θ*, and the second step shows that for all observations with a given **a**_**r**_, the likelihood depends only on deviation from the MLE and hence the likelihood functions are identical up to translation over the parameter space, with the function *f*( ·, **a**_**r**_ ) describing their common shape. Furthermore, the probability density of estimation error, 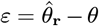, over all observations with a given **a**_**r**_ has the same shape as the corresponding likelihood functions. To be precise, the probability density function (PDF) is proportional to the likelihood for any one of the observations reflected and re-centred on zero (Efron & Hinkley, 1978, following Eq 2.3),

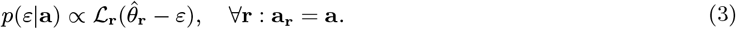

The shape of the PDF captured by 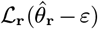 is invariant across all observations **r** satisfying **a**_**r**_ = **a**, even though its location in the parameter space, identified by the location of its peak 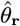, may differ (examples of likelihood functions with different *ε* but the same **a**_**r**_ are illustrated in Fig. 1).

**Figure 1.**
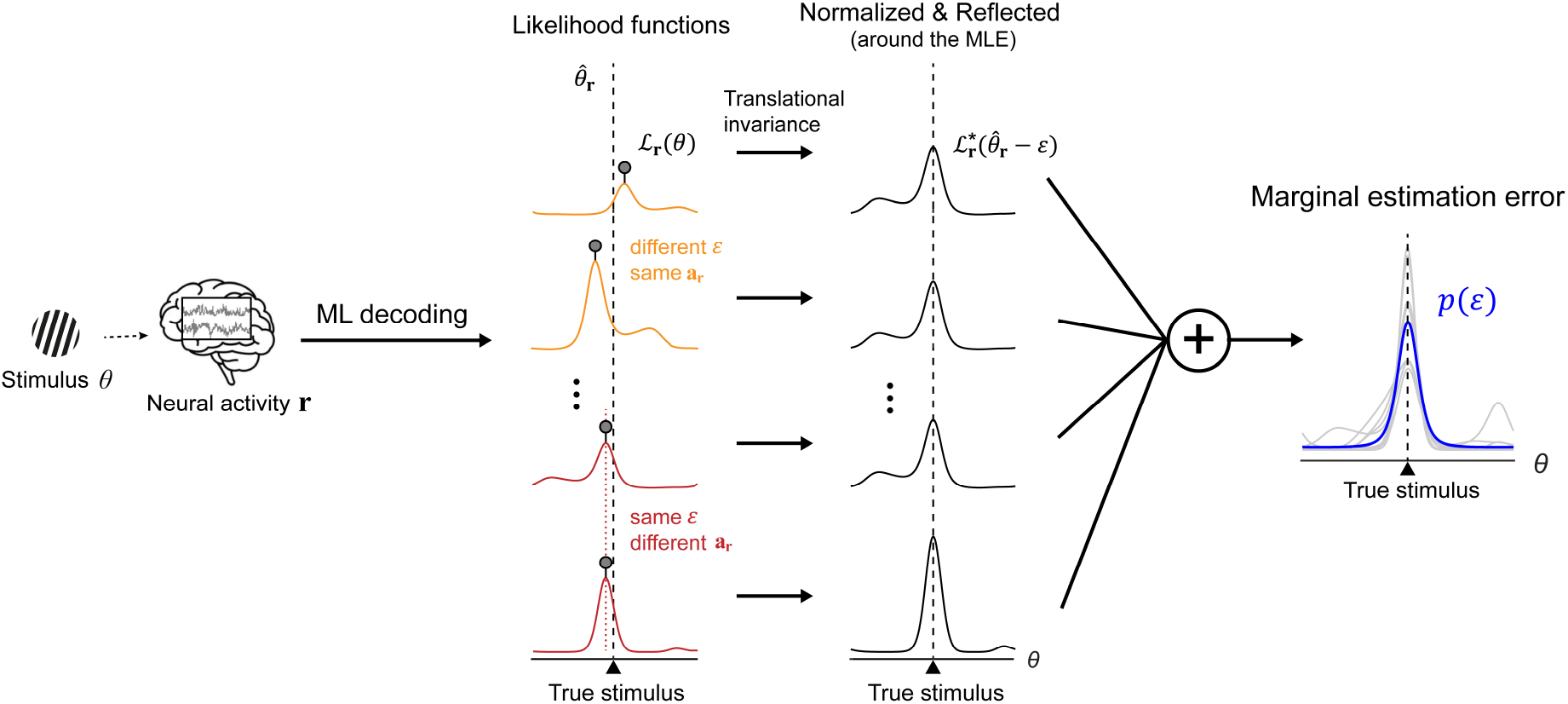
Marginal distribution of the estimation error in the MLE for translation-symmetric population codes. An example of maximum likelihood (ML) decoding: for a given stimulus (orientation *θ*), stochasticity in neural activity **r** leads to variability in likelihood width, shape and location of the peak (the 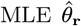). Among the different likelihood functions (left), the same estimation error *ε* (red curves; same 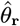 as indicated by the dotted red line) can be associated with different **a**_**r**_ (different shapes), whereas the same **a**_**r**_ (orange curves; same shape) can be associated with different values of *ε*. The marginal estimation error (blue curve; far right) is a mixture of the normalized, reflected (around the MLE) and re-centred likelihood components (grey curves; right).

Then, the marginal density of estimation error can be written as

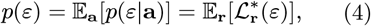

where 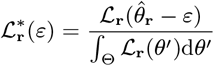

This says that the marginal error of 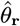 in estimating *θ* has a mixture distribution, with components consisting of the normalized, reflected (around the MLE) and re-centred likelihoods (Fig. 1). In subsequent sections we will see that *p*(*ε*) is generally heavy-tailed, and the tailedness is related to the information that can be extracted from ancillary statistics.

In behavioural estimation, ancillary statistics could form the basis of subjective uncertainty or confidence in a reported estimate. Here we consider two approaches to quantifying this “ancillary information”, one based on Fisher Information and statistical curvature – which yields some concise analytical results, though only for asymptotic conditions – and one based on Shannon’s Information Theory – which may be better suited for practical application to neural and behavioural data.

### 2.2 Quantifying ancillary information using Mutual Information

If we consider the parameter as well as the observation to be a random variable, the Mutual Information between them, denoted by MI(*θ*; **r**), quantifies how much an observation on average decreases uncertainty about the parameter. Mutual information is defined as MI(*x*; *y*) = *H*(*x*) − *H*(*x*|*y*), with differential entropy *H*(*x*) = −E_*x*_[log *p*(*x*)] and conditional differential entropy *H*(*x*|*y*) = −E_*x,y*_ [log *p*(*x*|*y*)].

Following Eq. 2, we can decompose MI(*θ*; **r**) into two parts: the information conveyed by the MLE 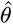 about the parameter *θ* and the information conveyed by the ancillary statistic **a**_**r**_ about the estimation error 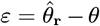,

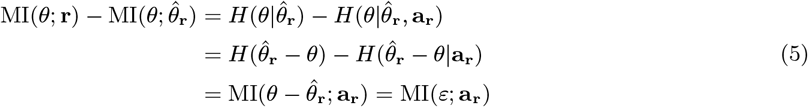

where we assume *p*(*θ*) constant, consistent with the translation-symmetric setting. Then we have

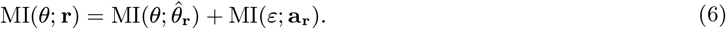

On the right-hand side (r.h.s.) in the above expression, the first term 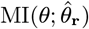 is a measure of estimation performance equal to the information the estimate 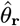 provides about the true parameter *θ* (Brunel & Nadal, 1998). The second term MI(*ε*; **a**_**r**_) measures the additional information about the estimation error *ε* (the accuracy of the MLE) provided by the ancillary statistic **a**_**r**_. A larger value of MI(*ε*; **a**_**r**_) indicates that the observations provide more information about uncertainty of the estimates. This mutual information can be expressed as the difference between the entropy of the marginal error in the MLE and the conditional entropy of the error given the population response

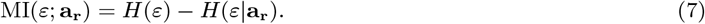

From Eq. 4 and Eq. 3, we have

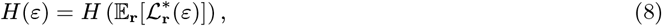

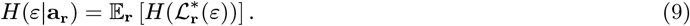

Thus, Eq. 7 can be rewritten in the form,

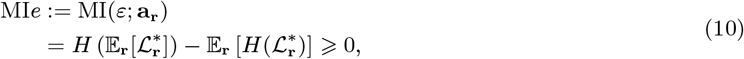

where we define the functional *H*(*f*) = − ∫_Θ_*f*(*θ*) log *f*(*θ*)d*θ*. In the following we will refer to this measure as MI*e*. Note that, because all the information about the error in the MLE is contained in **a**_**r**_, the measure can be equivalently defined as MI*e* = MI(*ε*; **a**_**r**_) or MI*e* = MI(*ε*; **r**).

Eq. 10 is a generalization of the Jensen–Shannon divergence (JSD), which measures the mean KL-divergence (a divergence based on Shannon information) of the mixture components to a mixture distribution (Burbea & Rao, 1982; J. Lin, 1991). In other words, the ancillary information grows with increasing diversity of the mixture components (i.e. the shapes of the likelihood functions). In general, a mixture distribution with more diverse components will have longer tails, so this result connects knowledge of uncertainty with heavy-tailed distributions of error, a point we explore further below.

### 2.3 Quantifying ancillary information using Fisher Information

Fisher Information is commonly used to characterize performance of an estimator, primarily because it provides the Cramér-Rao bound on estimation variability, which for an unbiased estimator 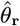 is,

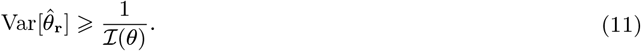

Here, the Expected FI, ℐ (*θ*), is a bound on average performance of the estimator; it has a sample-based counterpart in the Observed FI, *J*_*θ*_(**r**), the negative second derivative of the log likelihood, given a particular response **r**. The Observed FI can be evaluated at the MLE, in which case we write 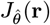, or at the true parameter, for which we write *J*_*θ*_(**r**); these are equivalent if the likelihood is Gaussian as *J*_*θ*_(**r**) is constant over the parameter space. By definition, the Expected FI is the expectation of the Observed FI evaluated at the true parameter, ℐ (*θ*) = E_**r**_[*J*_*θ*_(**r**)]. It can be approximated by the expected value of the Observed FI at the 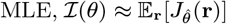, but they are not equal in general.

The inverse of the Observed FI evaluated at the MLE has been argued to provide the optimal estimator of the squared error of an estimate (Lindsay & Li, 1997). Efron and Hinkley (1978) suggested that the conditional variance 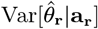 is more meaningful than 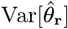 in assessing the precision of 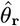 as an estimator of *θ*, with

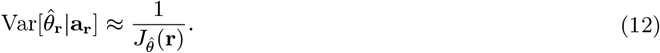

This approximation is exact for Gaussian likelihoods.

In a statistical model of the translation-type, when the MLE is insufficient, the Observed FI is an ancillary statistic that contains information about uncertainty of the MLE. Variation in the Observed FI across separate observations of the population implies variation in 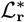 and hence the presence of ancillary information. If the Observed FI is sufficient to fully characterize the shape of the likelihood function, e.g. if the likelihoods have the same form differing only in location and scale (width), then the Observed FI is the ancillary complement to the MLE and contains all the information in the observation about reliability of the estimate.

R. A. Fisher (1925) described the “loss of [Fisher] information” when the 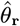 is substituted for the full observation **r**. This loss is equal to the FI provided by the ancillary statistic, **a**_**r**_. The Expected FI contained in the full observation is,

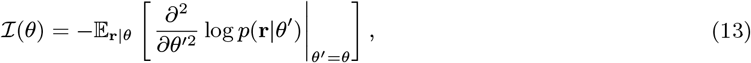

For a translation-type model, by Eq. 3, this is equivalent to,

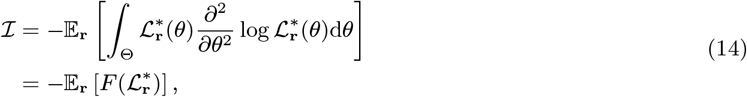

where we define the functional 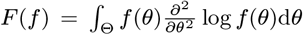. We have dropped the dependence on *θ* as the translational symmetry ensures the Expected FI is constant.

The Expected FI contained in 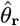 is,

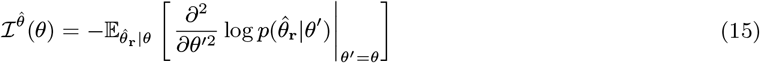

Following the same logic, for a translation-type model this is equivalent to,

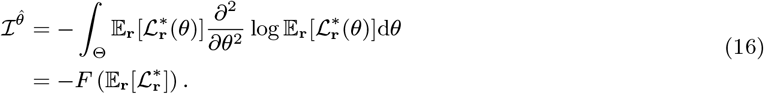

Giving,

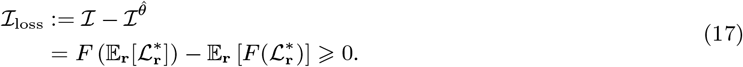

Like the mutual information measure of the previous section (Eq. 10), the information loss takes the form of a “Jensen difference”, a generalized diversity measure for probability distributions (Rao, 1982). In this case the functional *F*( ·) is based on Fisher instead of Shannon information, so Eq. 17 can be described as a generalized form of Jensen-Fisher divergence (JFD) (Sánchez-Moreno et al., 2012).

### 2.4 Relationship between the ancillary information measures

The information loss ℐ_loss_ (Eq. 17) and the mutual information MI*e* (Eq. 10) represent two alternative measures of ancillary information, i.e. the information contained in an observation about reliability of the MLE. Both measure the divergence between the distribution of estimation error and its component distributions (the recentred likelihood functions), with ℐ_loss_ (based on JFD) measuring the divergence in Fisher information and MI*e* (based on JSD) measuring it in terms of Shannon information. Thus the two measures differ on how they assess reliability of the MLE, mirroring the differences between the two conceptions of information: broadly speaking, for the FI-based ℐ_loss_, reliability is quantified as sensitivity to small changes in the true parameter, while for the entropy-based MI*e*, reliability reflects the probability of mistaking any one parameter value for another.

The two measures can be related via Stam’s inequality (Stam, 1959), which has the interpretation that of all distributions with a given Fisher information, i.e. 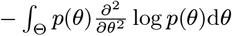, the Gaussian distribution has the minimum entropy (Wei & Stocker, 2016). It follows that, if we consider a model with a given ℐ_loss_ and marginal error distribution (fixing the first term in Eq. 10), the maximum mutual information MI*e* is obtained when the likelihoods (and errors conditioned on ancillary information) are Gaussian. Intuitively, the divergence in a set of Gaussian likelihoods is characterized equally well by Fisher information or entropy, so the ancillary information captured by ℐ_loss_ and MI*e* measures is the same, but if the likelihoods deviate from Gaussian, MI*e* may be sensitive to uncertainty about the true stimulus that ℐ_loss_ does not take into account.

### 2.5 Geometric view of statistical estimation and ancillary information

A geometric perspective on statistical estimation can further illuminate the relationship between estimation error and information about uncertainty. As a simple illustration, consider a game of boules (Fig. 2a), in which a heavy ball is thrown with the intention of landing as close as possible to a smaller target ball, the “jack”. Imagine that the heavy ball’s landing point has isotropic Gaussian variability centred on the jack, and consider an observer who knows the jack is at unit distance from the thrower but doesn’t know in which direction. Based on the heavy ball’s landing point, this observer’s maximum likelihood estimate of the direction of the jack from the thrower is the direction of the ball from the thrower. However, the reliability of this estimate increases the further the ball lands from the thrower: the distance is an ancillary complement.

**Figure 2.**
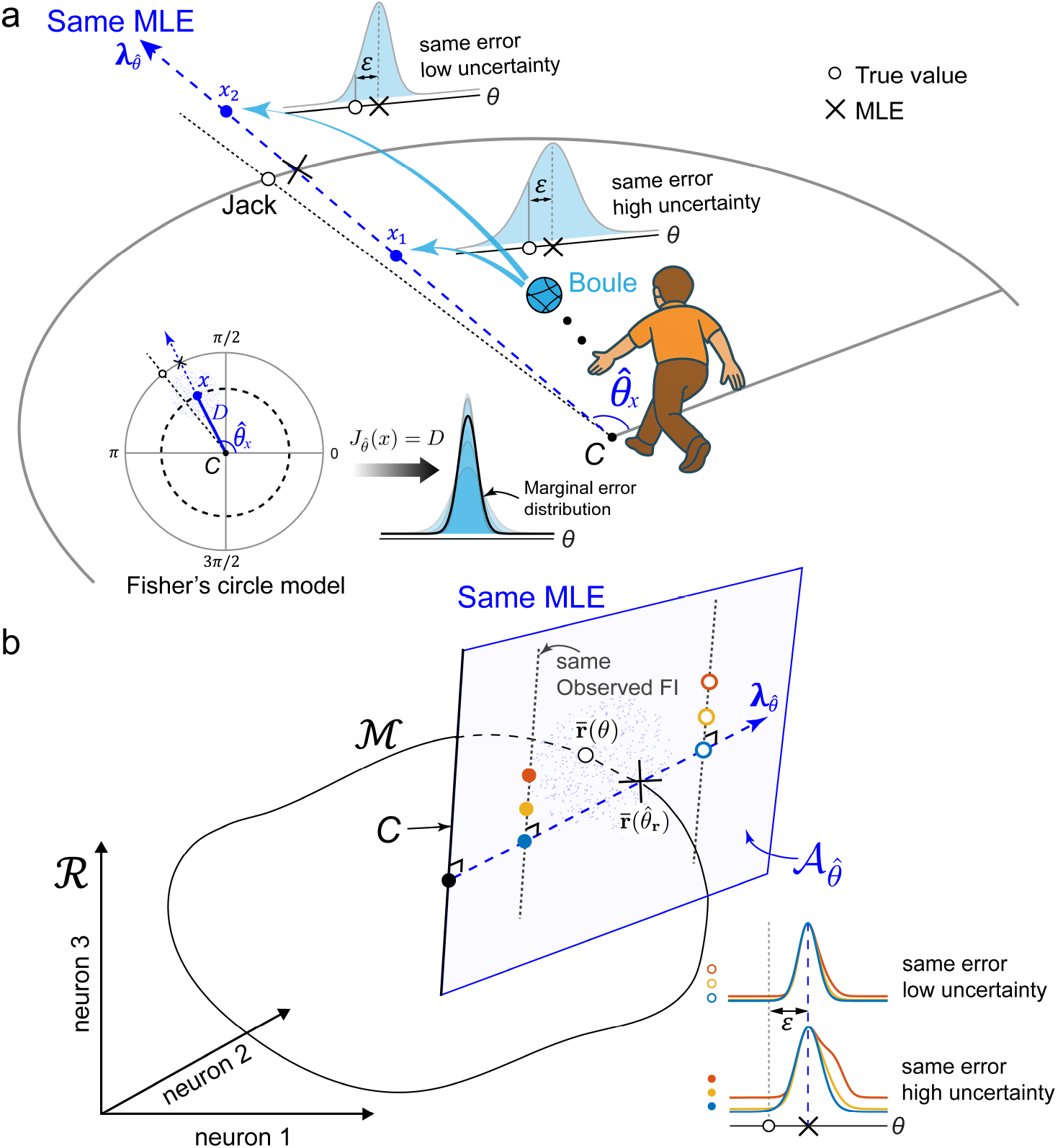
Geometric views of statistical inference for MLE. (a) In the boules example (see text) two different landing points (*x*_1_ and *x*_2_, blue dots) yield the same MLE 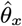 (black cross) of the direction of the jack, and thus the same angular error *ε* relative to the true direction (black unfilled dot), but the likelihoods differ in width, with the farther landing point having a narrower likelihood. The statistical model is equivalent to Fisher’s circle model (inset): likelihood functions are von Mises with concentration equal to the distance *D* from the centre of the unit circle. This is also the value of the Observed FI, which quantifies certainty in the estimate. The marginal error distribution is a scale mixture of von Mises with concentrations distributed identically to the distances. (b) Statistical inference based on high-dimensional neural activity **r** and exponential family noise: The mean response of *M* neurons forms a one-dimensional curved submanifold ℳ ⊂ ℛ. All observations **r** (blue dots) that yield the same 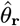 (black cross) lie in an (*M* − 1)-dimensional ancillary subspace 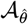. This ancillary subspace intersects with ℳ at 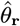. The critical point *C* is now an (*M ™* 2)-dimensional subspace containing responses that are equiprobable for all values of the stimulus *θ*. Within the ancillary subspace corresponding to a particular MLE, the Observed 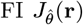 increases linearly with distance measured orthogonally from *C* in the direction of 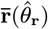, i.e. parallel to the vector 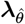 (unfilled dots correspond to observations with less uncertainty, as measured by Observed FI, than filled dots). Variation orthogonal to this line (e.g. differently coloured dots) corresponds to observations that are matched in estimation error and Observed FI, but differ in other aspects of the shape of the likelihood function (insets).

To make this more concrete, two different landing points (*x*_1_ and *x*_2_) are shown along with their corresponding likelihood functions, i.e. the probability of the landing point as a function of the direction of the jack, *p*(*x* | *θ*). The direction with maximum likelihood (the 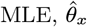) is the same for both of the illustrated landing points, so the estimation error is identical. However the likelihood functions differ in width, with the farther landing point having narrower likelihood, indicating less uncertainty about the jack’s direction: informally, this landing point is compatible with a smaller range of possible directions.

The statistical model here is equivalent to Fisher’s circle model (R. Fisher, 1955; described in Efron, 2022, Sec.4.6) for a circle with unit radius. Estimation certainty can be quantified by the Observed FI, which in this setting is exactly equal to the distance *D*,

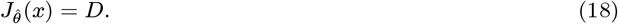

A landing point at the “critical point” *C*, at distance zero, would be equiprobable for all possible locations of the jack on the circle, and thus provide no information (uniform likelihood, 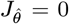). As distance of *x* from *C* increases, its projection onto the unit circle becomes increasingly sensitive to small changes in *θ*, so estimation certainty increases. The distance *D* from the critical point is ancillary, as it is statistically independent of the direction, and in this case it is the ancillary complement to the 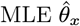, because together with the MLE it fully identifies the likelihood function, which is von Mises (circular normal) with circular mean 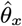 and concentration parameter equal to *D*.

The circle model is of the translation-type due to its rotational symmetry. As a result (Section 2.1) the distribution of the MLE can be predicted exactly from the form of the likelihood functions. Specifically, for a given distance *D* the MLE is von Mises distributed about the true location of the jack with concentration *D*, so the marginal error distribution is a scale mixture of von Mises with concentrations distributed identically to distances (Fig. 2a, inset). The mixture has heavier tails than a von Mises distribution due to the variation in width of the component distributions. In this case, the mixture distribution does not have a closed-form PDF; to further elucidate the relationship between ancillary information and tailedness, Section 2.6 describes a synthetic model with Gaussian likelihoods in which the marginal estimation error has a location-scale *t*-distribution.

The two-dimensional circle model provides a starting point for modelling estimation of a scalar stimulus value *θ* from a neural response **r** that lives in an *M* -dimensional space ℛ, where each dimension corresponds to the observed activity of one neuron (Fig. 2b). The circle (which consisted of the mean landing points for every different possible direction of the jack) can be generalized to a one-dimensional manifold ℳ of arbitrary shape embedded in ℛ consisting of the mean neural responses for every possible value of the stimulus, 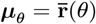.

Assuming the noise in neural activity belongs to the broad class of exponential family distributions, the statistical model takes the form of a one-parameter curved exponential family, with a well-characterized geometric interpretation (Efron, 1975). All observations **r** that yield the same 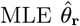 lie in a flat (*M* − 1)-dimensional hyperplane 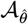 (Efron, 2022). This ancillary subspace intersects with ℳ at 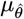 (Amari, 1982).

The critical point *C* now becomes an (*M* − 2)-dimensional critical subspace containing responses that are equiprobable for all values of the stimulus *θ*. Within the ancillary subspace corresponding to a particular MLE, the Observed FI 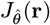 increases linearly with Euclidean distance measured orthogonally from *C* in the direction of 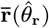, i.e. parallel to the vector labelled 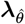. Variation in **r** within 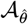 orthogonal to this line corresponds to observations that have the same MLE and Observed FI but differ in other aspects of the shape of the likelihood function.

The variation in Observed FI with distance from *C* is the cause of the information loss ℐ_loss_ described in the previous section. It arises because of the curvature of the statistical manifold defined by the curved exponential family. In the simplest case of unit isotropic Gaussian noise, the “statistical curvature” *γ*_*θ*_ is identical to the geometric curvature of the one-dimensional manifold ℳ in the response space ℛ. If the manifold were a straight line, the likelihood function and hence the Observed FI would be the same for all observations **r**. For translation-type models, this curvature directly determines the coefficient of variation in Observed FI (measured at the true parameter *θ*; see proof in Appendix)

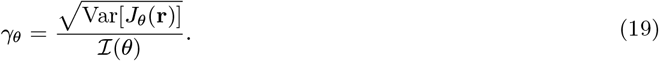

For other forms of noise distribution, the geometric curvature of ℳ would not sufficiently capture the variation in responses **r** about the mean response 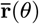 on ℳ. For an arbitrary noise distribution within the curved exponential family, the statistical curvature is instead measured on the manifold of *natural parameters* of the exponential family – in general, this manifold is non-Euclidean, with a metric that reflects the covariance structure of the noise (see Appendix for details).

The statistical curvature is related to the asymptotic information loss (Efron, 1975), the information loss in the limit in which estimation is based on many i.i.d. observations, by

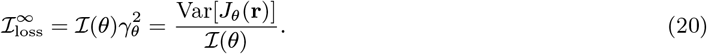

Note that ℐ (*θ*) = ℐand *γ*_*θ*_ = *γ* are constants due to the translational symmetry, so 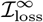 is invariant of *θ*. Our interest is in the information loss ℐ_loss_ as a measure of the ancillary information in a single observation of **r**(*θ*). Nevertheless, as we will see in the following sections, 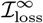 admits concise analytical expressions that provide intuition about the behaviour of ℐ_loss_ under different statistical models. However, the intuitive value of the geometric perspective for assessing ancillary information under non-Gaussian noise distributions is limited due to fact that the relevant curvature is measured on a non-Euclidean manifold. For clarity, we summarize the key statistical and information-theoretic quantities and their notation in Table. 1.

**Table 1.**
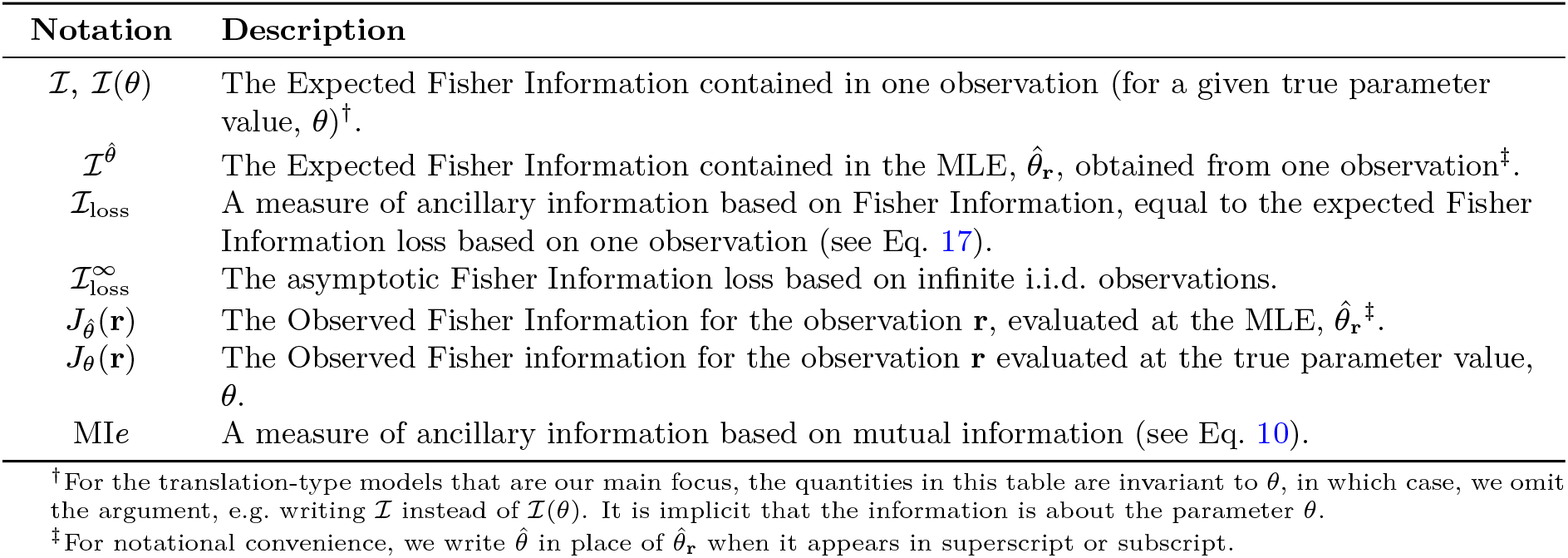
A summary of the key statistical and information-theoretic quantities.

### 2.6 Marginal estimation error as a mixture of normal distributions

To provide an intuitive illustration of how the distribution of estimation error is related to the loss of FI, we start with a simple example in which the marginal estimation error can be described as a mixture of Gaussian distributions.

Suppose that likelihoods ℒ_**r**_ are Gaussian distribution functions with the Observed FI drawn from a Gamma distribution (van den Berg et al., 2012), 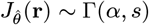, with 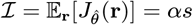, where *s* is the scale parameter and *α* is the shape parameter. The marginal distribution of estimation error is then expressed as a Gaussian scale mixture,

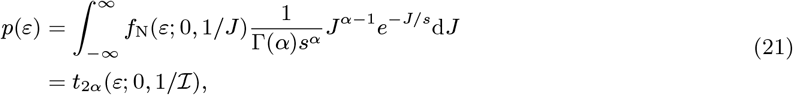

with 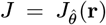. Here *f*_N_(*ε*; 0, 1*/J*) is the Gaussian PDF with zero mean and variance 1*/J*. The marginal error distribution is a location-scale *t*-distribution centred at 0, with degree of freedom *ν* = 2*α* and scale *τ* ^2^ = 1*/*(*αs*) = 1*/ℐ. ν* is also known as the normality parameter. Smaller values of *ν* correspond to heavier tails in the location-scale *t*-distribution, which approaches normality as *ν* → ∞.

By Eq. 17 and Eq. 20, ℐ_loss_ converges to 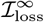 as *α* → ∞

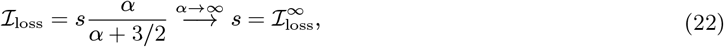

using 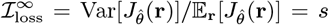. Since 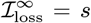, it follows that 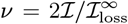. Therefore, for a fixed ℐ, increasing 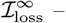 that is, greater variability in the Observed FI – results in heavier tails in the marginal error distribution.

For *ν* ≤ 2, the location-scale *t*-distribution is pathological since its variance is infinite or undefined. We next consider the more practical case of parameters in a circular space, where variances are always finite. Suppose the marginal error distribution is a mixture of von Mises (circular normal) distributions 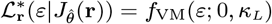, with 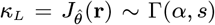, where *κ*_*L*_ is the likelihood concentration. Numerical simulations demonstrate that the shape of the marginal error distribution is again jointly determined by Expected FI, ℐ, and the loss of FI, ℐ_loss_. Specifically, increasing Expected FI leads to a narrower error distribution (Fig. 3a). Meanwhile, for a fixed Expected FI, larger values of 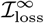 correspond to higher variability in the Observed FI, resulting in heavier tails in the marginal error distribution (Fig. 3b); in the limiting case where 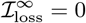, the error distribution reduces to a single von Mises distribution. Consistent with Eq. 22, ℐ_loss_ converges to 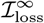 as *α* → ∞ (Fig. 3c). Both increases in ℐ_loss_ and 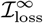 coincide with increases in MI*e* (Fig. 3d) across a substantial range; however, this relationship reverses for very large information loss.

**Figure 3.**
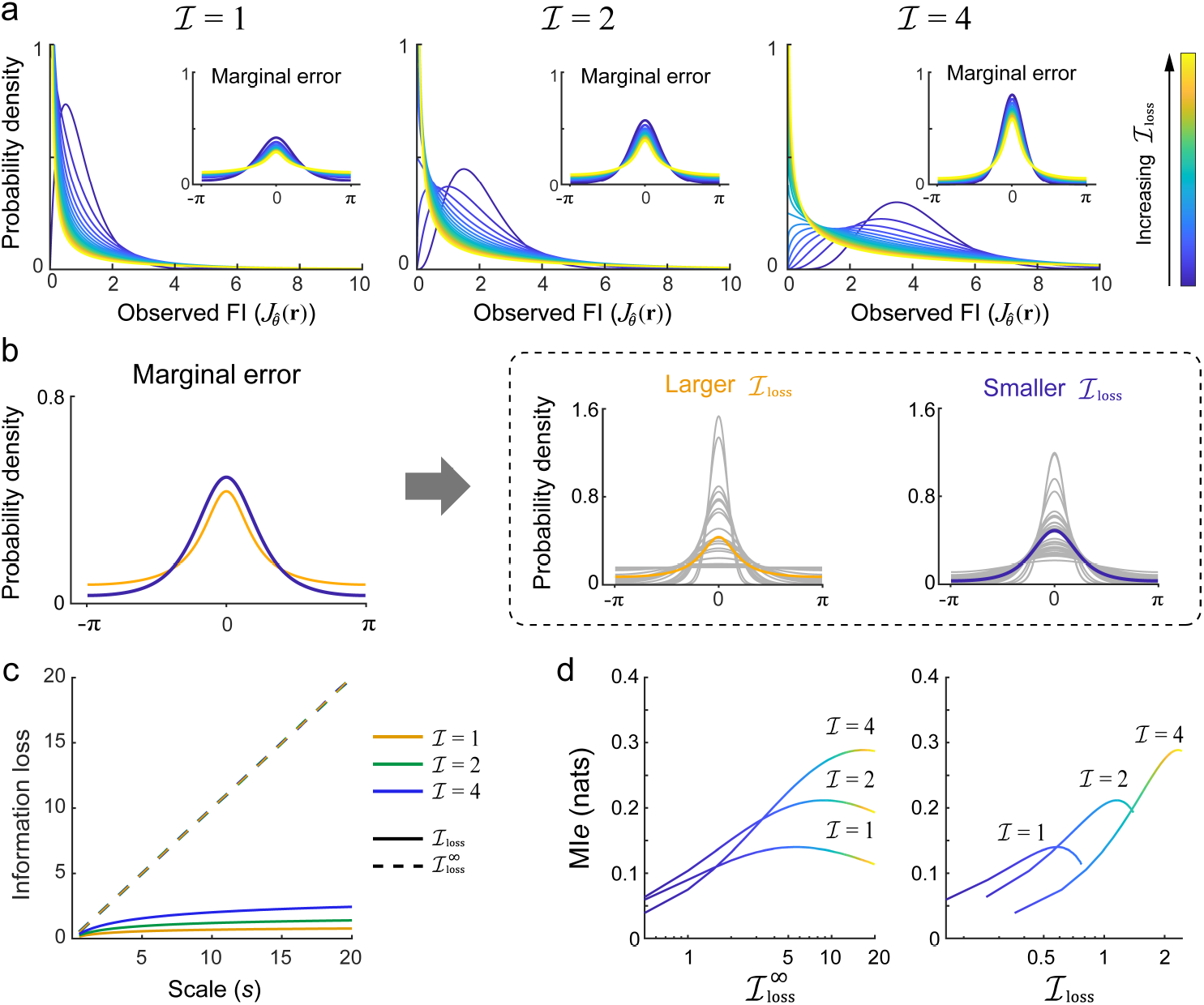
Marginal error as a mixture of circular normal (von Mises) distributions, with likelihood precision following a Gamma distribution. (a) The probability density of the Observed FI (Gamma distributed) and the corresponding marginal error distribution (insets), shown for fixed Expected FI ( ℐ = 1, 2, 4). Colours indicate different levels of 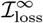. For a fixed ℐ, increasing 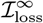 leads to heavier tails in the marginal error distribution. (b) Higher 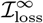 implies greater variability in the Observed FI, resulting in heavier tails in the marginal error distribution. (c) The information loss as a function of scale (*s*), which is related to the shape parameter *α* by *α* = ℐ */s*. For a fixed ℐ, the asymptotic loss (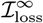, dashed lines) depends solely on *s* and increases linearly with it (corresponding to decreasing *α*). The information loss in a single observation (ℐ_loss_, solid lines) increases monotonically with *s*, plateaus at large *s* (with higher asymptote for larger ℐ), and converges to 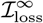 as *α* → ∞. (d) MI*e* as a function of ℐ_loss_ and 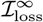 for fixed ℐ. Results obtained using Monte Carlo simulations with 10^7^ samples on a uniform grid of 201 points spanning *θ* ∈ [ − *π, π*). Unless otherwise indicated, these simulation settings are used throughout the paper.

### 2.7 Ancillary information in neural population codes

In this section we apply our measures of ancillary information to explicitly defined neural populations, to examine how different population characteristics influence the information available about uncertainty.

We assume a stimulus feature *θ* is encoded jointly in the population response **r** ≡ {*r*_1_, *r*_2_, …, *r*_*M*_ }of *M* neurons. The tuning functions **f** (*θ*) ≡ {*f*_1_(*θ*), *f*_2_(*θ*), …, *f*_*M*_ (*θ*) } along with a noise model *G*(**r, f** (*θ*)) define the probability of generating a particular population response,

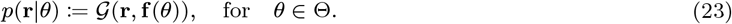

Consider tuning functions of the form *f*_*i*_(*θ*) = *f*_0_(*θ* − *φ*_*i*_), where the *φ*_*i*_ represent the preferred features of each neuron, which are uniformly distributed throughout the feature space Θ, and *f*_0_(· ) determines the shape of tuning function. In the continuum limit of large *M*, such a statistical model is of the translation-type with *θ* as a location parameter, so we can apply the results obtained in the previous sections.

Assuming neurons’ responses are independent conditioned on *θ*, the likelihood can be written as

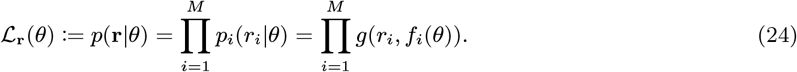

If we further assume that the distribution of each neuron’s response conditioned on stimulus *θ* has the form of an exponential family

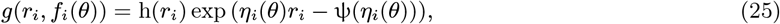

with *η*_*i*_(*θ*) = *η*(*f*_*i*_(*θ*)). Then the likelihood function is given by

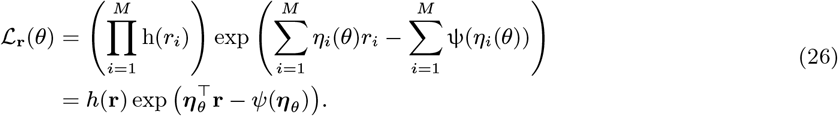

with ***η***_*θ*_ = {*η*(*f*_1_(*θ*)), *η*(*f*_2_(*θ*)), …, *η*(*f*_*M*_ (*θ*)) } . Since *η*(· ) is generally a nonlinear function of *θ*, ℒ_**r**_(*θ*) has the form of a curved exponential family. The curvature, which is determined by the combination of tuning functions and noise distribution, leads to variation in likelihood width and shape, producing ancillary information about uncertainty and heavy tails in the marginal error distribution (Fig. 4a).

**Figure 4.**
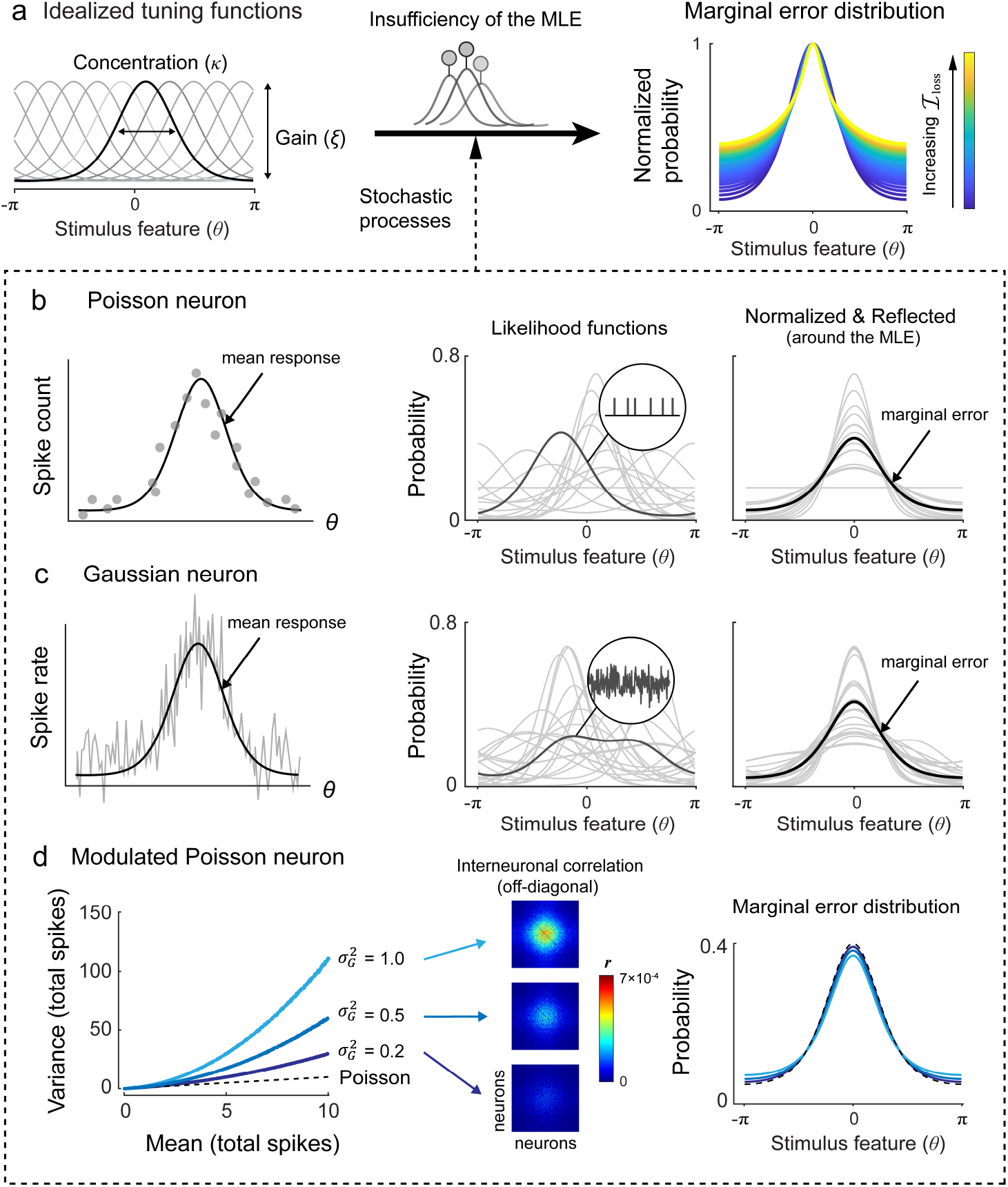
The neural population coding framework. (a) Assuming idealized identically shaped tuning functions densely and evenly distributed over a circular stimulus space, noise in the neural response leads to insufficiency of the MLE, manifested in variation of likelihood width and shape and yielding heavy tails in the marginal error distribution. With a fixed value of Expected FI, for small ℐ_loss_, the marginal error distribution is approximately a circular normal distribution (dark blue curves). As ℐ_loss_ increases, the error distribution deviates more from normality (light yellow curves). Error distributions are normalized by peak probability. (b) In the Poisson noise model, spike counts are discrete and non-negative, resulting in normal (von Mises) likelihoods. (c) In the Gaussian noise model, response amplitudes are continuous and can include negative responses, and likelihoods are non-normal and may be multimodal. (d) For the modulated Poisson model with overdispersed Poisson firing, larger 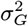 leads to greater overdispersion in the neural response, resulting in heavier tails in the marginal error distribution and stronger interneuronal correlations. The interneuronal correlation matrix is shown with each off-diagonal entry representing the activity correlation between a pair of neurons; diagonal elements (autocorrelations) are omitted for illustrative purpose. Simulations use parameters of *κ* = 1, *ξ* = 2 for the (modulated) Poisson models, and equivalents for the Gaussian model based on the variance-stabilizing reparametrization. Results obtained with a population of *M* = 200 simulated neurons; the same setting is used throughout unless otherwise indicated.

We will make the common assumption of von Mises (circular normal) tuning functions,

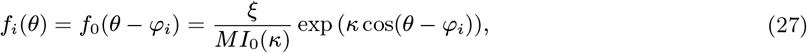

with tuning concentration (inverse width) *κ* and population gain *ξ. I*_*n*_( ·) is the modified Bessel function of the first kind.

For certain noise models (Fig. 4b-d; described in detail below), the form of the likelihood function can be specified explicitly, and the analytic expressions of the expectation and variance of the Observed FI, and thus 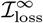, can be straightforwardly obtained from their curved exponential forms, as summarized in Table. 2 (see Appendix for derivations).

Assuming uniform coverage of neurons, 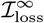 solely depends on tuning concentration *κ* in the Poisson and Gaussian noise models, whereas it depends on both gain *ξ* and modulation parameter 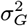 in the modulated Poisson model. For all the models, 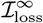 monotonically increases with *κ* (increasingly narrow tuning), and 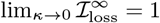. Fig. S2 shows how these quantities change depending on *κ* and *ξ* in the Poisson, Gaussian and modulated Poisson noise models. When Expected FI is held constant, increasing *κ* corresponds to decreasing *ξ*. Note that our results are obtained by specifying the expected summed activity (*ξ*) and tuning concentration (*κ*), while assuming the number of neurons is always sufficient to densely tile the stimulus space with tuning functions; for a fixed and finite number of neurons, increasing tuning concentration *κ* would also have the effect of decreasing coverage of the stimulus space, the effects of which are not considered here.

#### 2.7.1 Ancillary information depends on the noise model

##### Poisson noise

The standard Poisson noise model assumes that neural responses encoding stimulus feature *θ* are Poisson distributed (Ma et al., 2006; Pouget et al., 2003; Seung & Sompolinsky, 1993), *r*_*i*_ | *θ* ∼ *P*(*f*_*i*_(*θ*)), where neural spike count is discrete and non-negative (Fig. 4b). Assuming independent noise, the joint likelihood function is given by

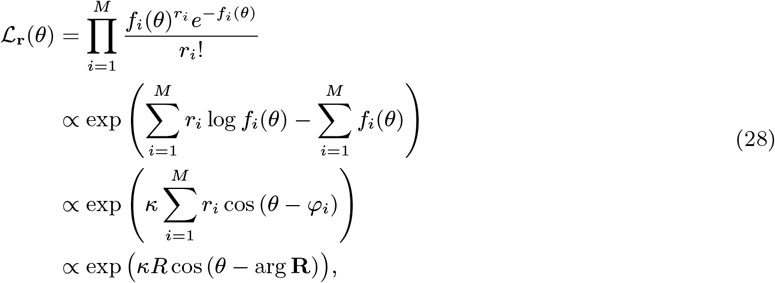

with 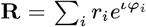 (the resultant vector) and *R* = |**R**| (the resultant vector length), where we have used that 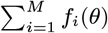 is approximately constant for large *M*.

Associating each spike *j* with the preferred value of the neuron that produced it *φ*_(*j*)_, we can also write 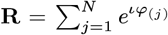, i.e. the resultant vector of spike values. (Note that *j* takes values from 1 to *N* = ∑ _*i*_*r*_*i*_, the total number of spikes.) **R** is a sufficient statistic for *θ* and can be decomposed into the MLE, arg **R**, which is the circular mean of the spike values, and the resultant length *R* which is an ancillary complement containing all the information in the neural response about the precision of the MLE. For a given resultant length, the likelihood function has the shape of a von Mises with concentration *κR*, which is also the value of the Observed 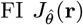, and the MLE is von Mises distributed with the same concentration centred on *θ*. Thus the marginal error distribution is a scale mixture of von Mises with concentrations having the same distribution as *R*, scaled by *κ*. The distribution of *R* does not have a closed-form representation in general.

##### Gaussian approximation

Many studies choose to simplify analysis by assuming Gaussian noise in population responses (Ma et al., 2006; Schneegans & Bays, 2017; Wei & Woodford, 2025; Wilke & Eurich, 2002; Wu et al., 2002). Although the Gaussian assumption implies negative responses that are biologically implausible, it remains a mathematically powerful tool for handling continuous variables and covariance, and models based on it have provided good fits to behavioural error patterns (Schneegans & Bays, 2017; Schurgin et al., 2020; Wei & Woodford, 2025). In the Gaussian noise model, neural activity is corrupted by additive (or “flat”) Gaussian noise, giving neural responses *r*_*i*_ | *θ* ∼ *N*(*f*_*i*_(*θ*), *σ*^2^). We set *σ*^2^ = *ξ/M* so that variance is equal to the mean population activity, approximating Poisson noise. The likelihood function is given by

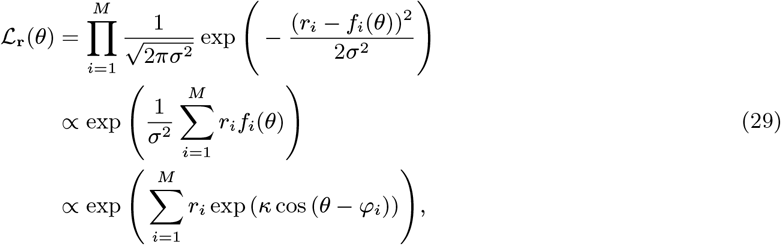

which is in general not proportional to a von Mises (Fig. 4c) nor sufficiently characterized by the Observed FI, which is not an ancillary complement in this case. In fact, as the log likelihood is equal (up to an additive constant) to a sum of von Mises functions weighted by the responses, and finite mixtures of von Mises distributions are identifiable (Fraser et al., 1981), it follows that the number of distinct identifiable log likelihood functions grows without bound with *M*. Because a sufficient statistic must identify a unique log likelihood (up to an additive constant), this model does not admit a sufficient statistic that remains finite as *M* → ∞. As a corollary there is no finite ancillary complement, analogous to the resultant length in the Poisson model, as *M* → ∞. In simple terms, information about uncertainty of the MLE is distributed throughout the whole pattern of activity in this model, meaning that it cannot be effectively summarized in a statistic of smaller dimension.

Let *κ* and *ξ* denote the concentration and gain parameters for the Poisson noise model and correspondingly *κ*^*′*^ and *ξ*^*′*^ for the Gaussian noise model. To better equate the Poisson noise model to the flat Gaussian noise model, we apply a variance-stabilizing transformation to the Poisson neurons’ responses *r*_*i*_, yielding the following reparametrization (see Appendix for derivations):

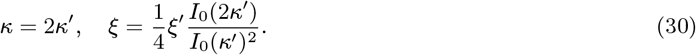

This reparametrization yields nearly identical error patterns between the two models (Fig. 5a), as noted in previous work (Tomić & Bays, 2024b).

**Figure 5.**
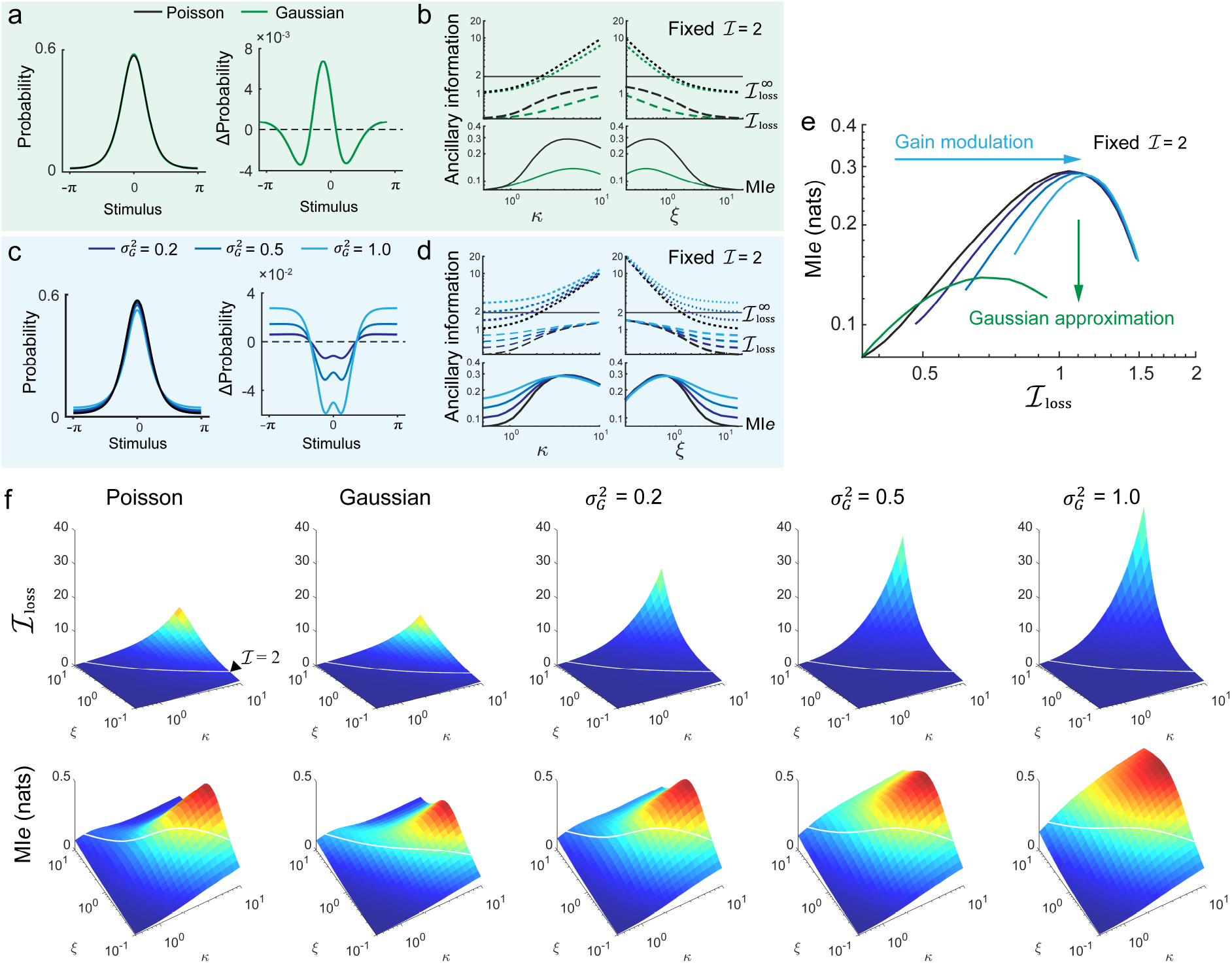
Ancillary information in different noise models. (a) A Gaussian approximation to Poisson variability (green curves) based on a variance-stabilizing transformation yields similar error distribution pattern to the standard Poisson model (black curves). (b) Ancillary information as a function of parameters *κ* and *ξ*, with the Expected FI held fixed (ℐ = 2). Under this constraint, increasing *κ* is accompanied by decreasing *ξ*. With matched parameters, the Gaussian approximation has the same Expected FI as the Poisson but less ancillary information in terms of MI*e* (solid curves), ℐ_loss_ (dashed curves) and asymptotically as 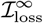 (dotted curves). (c-d) Similar to (a-b) but with varying Poisson modulation parameter 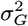. Modulation increases ancillary information (e) The relationship between MI*e* and ℐ_loss_ in different noise models, with fixed Expected FI ( ℐ = 2). The standard Poisson model corresponds to the black line. (f) Ancillary information dependent on parameters *κ* and *ξ* in terms of ℐ_loss_ (upper panel) and MI*e* (lower panel). White contour lines indicate ℐ= 2.

Plugging Eq. 30 into the expressions of E_**r**_[*J*_*θ*_(**r**)] and Var_**r**_[*J*_*θ*_(**r**)] for the Poisson model in Table. 2, we have the expectation and variance of Observed FI for the Poisson model in terms of the parameters of the matching Gaussian model as

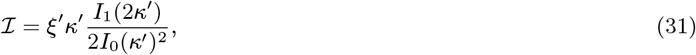

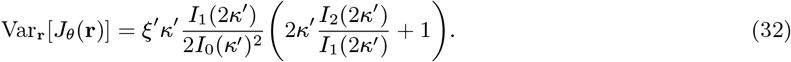

**Table 2.**
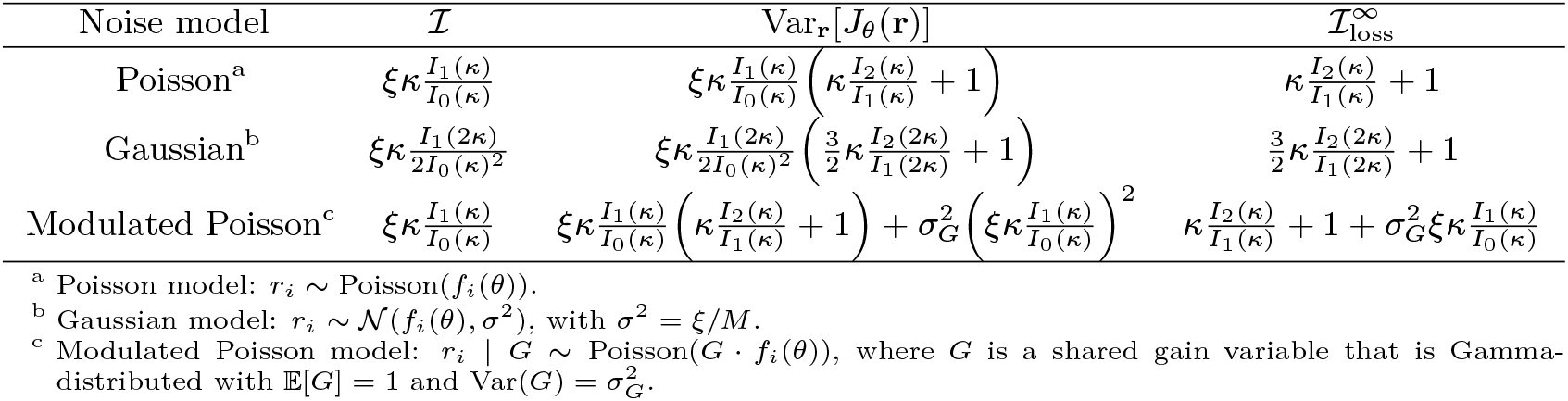
FI and the asymptotic loss of FI in the MLE for some statistical models.

| Noise model | $\mathcal{I}$ | $\text{Var}_{\mathbf{r}}[J_{\theta}(\mathbf{r})]$ | $\mathcal{I}_{\text{loss}}^{\infty}$ |
| --- | --- | --- | --- |
| Poisson<sup>a</sup> | $\xi \kappa \frac{I_1(\kappa)}{I_0(\kappa)}$ | $\xi \kappa \frac{I_1(\kappa)}{I_0(\kappa)} \left( \kappa \frac{I_2(\kappa)}{I_1(\kappa)} + 1 \right)$ | $\kappa \frac{I_2(\kappa)}{I_1(\kappa)} + 1$ |
| Gaussian<sup>b</sup> | $\xi \kappa \frac{I_1(2\kappa)}{2I_0(\kappa)^2}$ | $\xi \kappa \frac{I_1(2\kappa)}{2I_0(\kappa)^2} \left( \frac{3}{2} \kappa \frac{I_2(2\kappa)}{I_1(2\kappa)} + 1 \right)$ | $\frac{3}{2} \kappa \frac{I_2(2\kappa)}{I_1(2\kappa)} + 1$ |
| Modulated Poisson<sup>c</sup> | $\xi \kappa \frac{I_1(\kappa)}{I_0(\kappa)}$ | $\xi \kappa \frac{I_1(\kappa)}{I_0(\kappa)} \left( \kappa \frac{I_2(\kappa)}{I_1(\kappa)} + 1 \right) + \sigma_G^2 \left( \xi \kappa \frac{I_1(\kappa)}{I_0(\kappa)} \right)^2$ | $\kappa \frac{I_2(\kappa)}{I_1(\kappa)} + 1 + \sigma_G^2 \xi \kappa \frac{I_1(\kappa)}{I_0(\kappa)}$ |

<sup>a</sup> Poisson model: $r_i \sim \text{Poisson}(f_i(\theta))$ .
<sup>b</sup> Gaussian model: $r_i \sim \mathcal{N}(f_i(\theta), \sigma^2)$ , with $\sigma^2 = \xi/M$ .
<sup>c</sup> Modulated Poisson model: $r_i | G \sim \text{Poisson}(G \cdot f_i(\theta))$ , where $G$ is a shared gain variable that is Gamma-distributed with $\mathbb{E}[G] = 1$ and $\text{Var}(G) = \sigma_G^2$ .

Note that the r.h.s. of Eq. 31 has the exact form of ℐ in the Gaussian model, indicating that the variance-stabilizing reparametrization equates the Expected FI between the two models. Thus the difference in 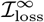 is solely determined by the difference in Var_**r**_[*J*_*θ*_(**r**)], with the Poisson model exhibiting higher Var_**r**_[*J*_*θ*_(**r**)] by

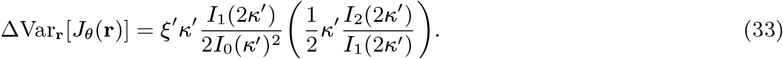

Consequently, the Poisson model exhibits higher 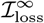 than the Gaussian model by

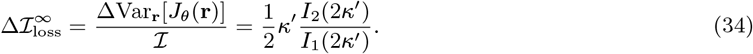

This says that approximating Poisson noise with Gaussian noise leads to an underestimation of 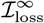 for *κ >* 0 that increases with *κ* without bound. Numerical estimation confirms a similar systematic underestimation of ancillary information measured both as ℐ_loss_ and MI*e* (Fig. 5b), with this underestimation also observed over the full parameter space (Fig. 5f).

Because the Gaussian noise model has non-normal likelihood functions that are not fully identified by the Observed FI (Eq. 29), the approximation of conditional variance by 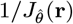 (Eq. 12) loses information contained in the ancillary statistic. This marks a difference from the Poisson noise model. Approximating Poisson activity as Gaussian underestimates the available ancillary information, and because the Expected FI (the total information contained in **r**) is preserved, also exaggerates the Fisher information carried by the 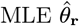. Both these discrepancies grow with increasing *κ*.

##### Gain modulation

The standard Poisson model predicts variance equal to the mean (Fano factor = 1) for the neural spike count. However, this assumption does not hold exactly for biological neurons, with overdispersion (Fano factor *>* 1) widely observed in electrophysiological recordings (Churchland et al., 2010; Graf et al., 2011; Ponce-Alvarez et al., 2013; Taouali et al., 2016; Zohary et al., 1994). The modulated Poisson model accounts for overdispersion by modelling spiking as a doubly stochastic process, where Poisson spike rate is modulated by a multiplicative stochastic gain *G* (Goris et al., 2014; Pillow & Scott, 2012; Taouali et al., 2016). Specifically, the mean response of the *i*th neuron to stimulus *θ* is 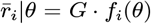, where *G* is a Gamma-distributed random variable, with shape parameter 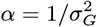 and scale parameter 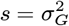, such that E[*G*] = 1 and 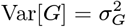. In this case, the variance of the response given *θ* can be described as

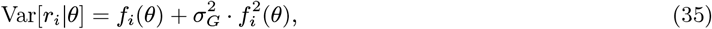

giving a Fano factor of

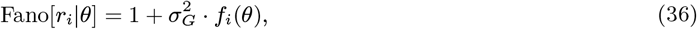

which increases linearly with 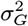. As 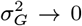, the modulated Poisson model reduces to the standard Poisson model (see Fig. 4d).

Given the evidence for shared neural variability (I.-C. Lin et al., 2015), we are particularly interested in the case where gain fluctuations are shared across neurons. The likelihood ℒ_**r**_(*θ*) is obtained by marginalizing over the shared modulation variable *G*,

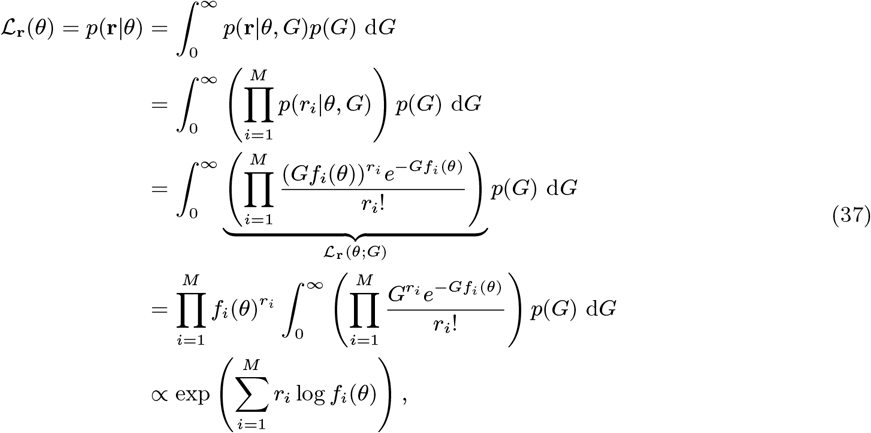

where ℒ_**r**_ (*θ* ; *G*) is the likelihood function of the standard Poisson model with population gain *G* · *ξ*, and we have again made use of 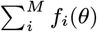 being approximately constant for large *M*, as well as that spike counts are conditionally independent for a given modulated gain *G*. The last line in the above expression indicates that gain modulation does not affect the likelihood function given a particular response **r**, which is von Mises and characterized by the resultant vector **R** exactly as for the standard Poisson (Eq. 28). Differences from the Poisson model therefore arise solely from the effects of modulation on the distribution of resultant length *R*, which in turn determine the distribution of Observed 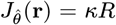 and the distribution of von Mises concentrations in the scale mixture that comprises the marginal error distribution.

Note that gain modulation introduces interneuronal correlation without adding explicit synchronization between neurons. Based on the properties of the multivariate Poisson-Gamma mixture model (Winkelmann, 2008, Section 7.1.3), the covariance matrix of **r** is given by

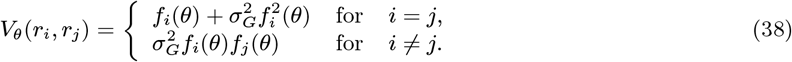

Interneuronal correlations as well as individual firing rates can carry information about a stimulus variable (Ponce-Alvarez et al., 2013), however, as shown in Table. 2, given parameters *κ* and *ξ*, the modulated Poisson model yields the same Expected FI as the standard Poisson model. This indicates that shared gain modulation, despite introducing interneuronal correlations, does not change the Expected FI, the average information contained in the neural response **r**. However, gain modulation systematically increases 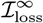 compared to the standard Poisson model, with an additional asymptotic loss that increases linearly with the modulation parameter 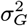,

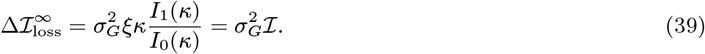

Effects on both ℐ_loss_ and MI*e* are qualitatively consistent with those for 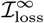 (Fig. 5d), indicating that ancillary information increases monotonically with strength of modulation. Since the total information (the Expected FI) is independent of modulation strength, the increase in information loss implies a decrease in the information carried by the 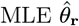.

##### Dissociation between information loss and Mie

Notably, there is a systematic dissociation between the FI- and entropy-based ancillary information measures. Fig. 5f shows that the two measures have distinct dependences on the tuning concentration (*κ*) and the population gain (*ξ*): ℐ_loss_ increases without bound as either parameter increases, whereas MI*e* exhibits an inverted-U pattern with *ξ* for *κ* constant and non-zero.

The dissociation between ℐ_loss_ and MI*e* is also evident when the Expected FI is held fixed at a non-zero value (e.g., ℐ = 2 in Figs. 5bd&e, also shown as white contour lines in Fig. 5f). With ℐ held constant, increasing *κ* is accompanied by decreasing *ξ*, and ℐ_loss_ grows monotonically towards an asymptote equal to ℐ as *κ* → ∞ (correspondingly *ξ* → 0), whereas MI*e* first increases and then decreases (Figs. 5b&d). Since ℐ_loss_ is monotonic in *κ*, it translates to an inverted-U dependence between MI*e* and ℐ_loss_ (Fig. 5e).

This non-monotonic relationship reflects the fact that MI*e* tends to zero in both limiting regimes. As ℐ_loss_ → 0, likelihoods and uncertainty become nearly identical across observations and thus carry little information about the estimation error, so MI*e →* 0. Conversely, as ℐ_loss_ → ℐ (corresponding to *κ* → ∞ and *ξ* → 0), likelihoods strongly diverge, with the majority of observations having very broad likelihood functions and very low certainty and a small minority of very narrow likelihoods with high certainty. The former observations dominate the mutual information measure so MI*e* again tends to zero.

Finally, although the inverted-U dependence between ℐ_loss_ and MI*e* is preserved across models, the magnitude of MI*e* can differ substantially even when matched for both ℐ and ℐ_loss_. Relative to the standard Poisson model, the Gaussian approximation has strongly reduced MI*e* at large ℐ_loss_, whereas gain modulation displays the opposite pattern, with the strongest reduction in MI*e* at small ℐ_loss_ (Fig. 5e). The entropy-based measure of ancillary information MI*e* thus allows further differentiation between competing models that are indistinguishable based on the combination of FI and loss of FI.

#### 2.7.2 Effects of external noise versus weakening activity

As population tuning is relatively static, changes in the quality of stimulus representation in a neural population will typically reflect either a change in activity amplitude (*ξ*) or additional variability accrued in the represented value itself. Examples of the latter, which we will call *external noise*, could include variability in sensory transduction or neural transmission that corrupts the signal before it is encoded in the population of interest (Faisal et al., 2008; Meister & Berry, 1999; Pelli, 1991), or accumulation of random error induced by recurrent connections that sustain a signal over time (Burak & Fiete, 2012; Kim et al., 2017; Schneegans & Bays, 2018; Tomić & Bays, 2024a; Wimmer et al., 2014; Wolff et al., 2020). Intuitively, the addition of external noise increases estimation uncertainty without contributing information about the uncertainty, and so it may affect ancillary information measures differently from a reduction in activity amplitude. Note that this contrast is the same one sometimes described as external versus *internal noise*, although strictly it is the internal signal-to-noise ratio (SNR) that changes.

To investigate how external noise affects information in a model population, we assume that the encoded value 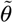 differs from the true stimulus value *θ* by addition of a Gaussian random variable with zero mean and variance 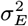,

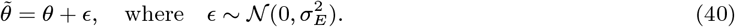

Let ℒ_**r**_ and 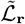 denote the likelihood without and with adding external noise. The resulting likelihood is given by convolving ℒ_**r**_ with a Gaussian kernel *K*_*σ*_

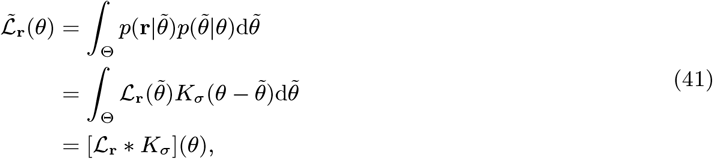

where 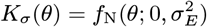 is a Gaussian distribution with zero mean and variance 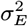.

Next, we consider a simple case where the marginal error is a mixture of Gaussian likelihoods to understand how external noise affects estimation performance of the MLE. The convolution preserves the location of peak likelihood (the MLE) but increases the likelihood width as

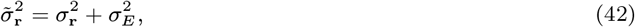

where 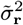 and 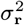 denote the variance of the normalized likelihoods 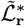 and 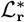, respectively. The marginal variance 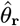 is given by

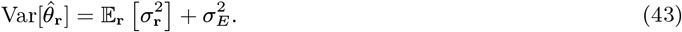

Here, 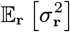 is the marginal variance without the external noise. This indicates that adding external noise increases the marginal variance of 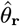, manifested in the increasing width of the marginal error distribution (Fig. 6a). Let 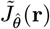 and 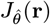 denote the Observed FI given 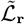 and ℒ_**r**_, respectively. Following Eq. 42, we have

**Figure 6.**
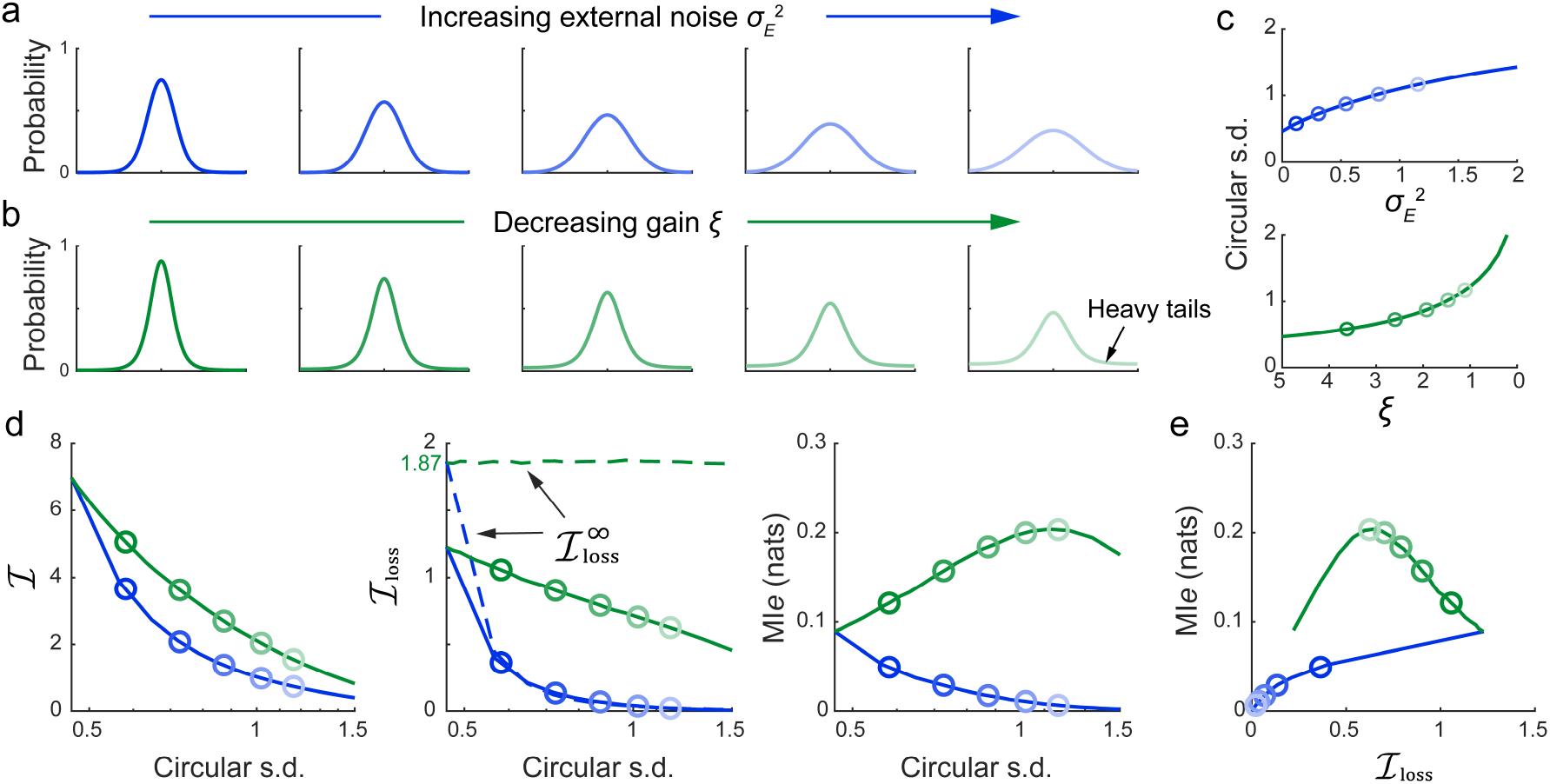
Effects of increasing external noise versus weakening activity. (a) Increasing external noise 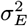 (with fixed *κ* = 2 and *ξ* = 5) and (b) decreasing population gain *ξ* (with fixed *κ* = 2) both produce increased error variability (vertically aligned panels are matched for circular s.d.). Decreasing *ξ* leads to heavy tails in the marginal error distribution. (c) Changes in the error variability (circular s.d.) dependent on 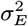 and *ξ*. (d) Ancillary information (i.e. ℐ_loss_, 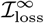 and MI*e*) as a function of changes to circular s.d. induced by external noise (blue curves) or weakening activity (green curves). Dashed curves represent the values of 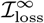. The theoretical value of 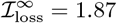 is calculated as *κI*_2_(*κ*)*/I*_1_(*κ*) + 1 (Table. 2, the Poisson model). (e) The relationship between the two ancillary information measures with respect to changes in external noise (increasing 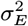, blue curve) or weakening activity (decreasing *ξ*, green curve). All simulations are based on the standard Poisson noise model. Coloured open circles in (c–e) correspond to the error distributions shown in (a,b).

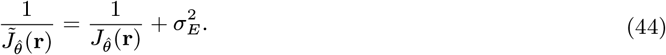

It is easy to see that, for positive 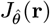, adding external noise reduces the Expected FI (Fig. 6c)

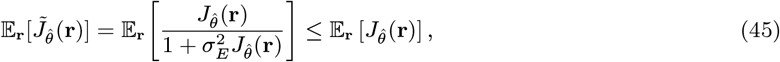

while equality holds for 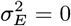.

We now extend the formulation to the circular case by considering convolution of von Mises likelihoods (as in e.g., the Poisson noise model) with a wrapped normal distribution *K*_*σ*_(*θ*) = *f*_WN_(*θ*; 0, *σ*_*E*_) with zero mean and circular standard deviation (s.d.) *σ*_*E*_ . Note that, for large mean resultant length, the von Mises likelihood can be approximated by a wrapped normal distribution. So convolution with a wrapped normal distribution with circular s.d. *σ*_*E*_ can be closely approximated by scaling the resultant length of the von Mises likelihoods by the resultant length of the wrapped normal 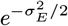 (Wernicke et al., 2026),

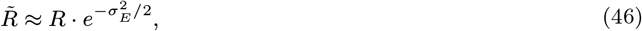

with 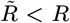 for *σ*_*E*_ *>* 0. As the Observed FI is monotonic in the resultant length, it follows that all Observed FIs and the Expected FI will decrease monotonically with increasing external noise.

Accordingly, the circular s.d. of 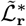 increases with external noise

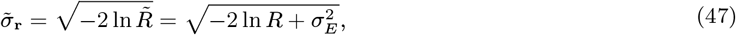

or equivalently, 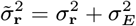, which preserves the additive property of the linear case (Eq. 42).

Numerical simulations confirmed that, given fixed *κ* and *ξ*, increasing external noise (*σ*_*E*_ ) leads to greater error variability, indicated by the increasing circular s.d. in the marginal error distribution (Fig. 6a & c). This effect is similar to that of decreasing the population gain *ξ*, when *κ* is fixed (Fig. 6b & c). Both manipulations monotonically reduce the Expected FI (ℐ). The asymptotic information loss, 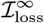 is independent of *ξ* (Table. 2, the Poisson model), so weakening activity does not affect the asymptotic loss, while adding external noise significantly reduces it, with greater reduction for larger error variability. The actual information loss (ℐ_loss_) is reduced by both manipulations, but external noise causes greater reduction for a matched effect on error variability.

The largest contrast is observed for the entropy-based measure of ancillary information MI*e*, which also decreases monotonically with increasing external noise 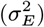, but follows an inverted-U pattern as a result of weakening activity (decreasing *ξ*), such that the two manipulations can have opposite affects on MI*e* for small matched increases in error variability (Fig. 6d). Similar patterns are observed in MI*e* even when matching for ℐ_loss_ (Fig. 6e).

## 3 Discussion

We have presented a new framework relating principles of neural coding to information about uncertainty and non-Gaussianity in sensory estimates. We showed that ancillary information – by which we mean, the information about a stimulus contained in population activity additional to that expressed in the MLE – manifests as variability in the width and shape of the likelihood function. This variability naturally gives rise to non-Gaussianity of error distributions based on the MLE, even when individual likelihoods are Gaussian functions, and provides a basis for uncertainty to be informative about estimation error, a property known as metacognitive sensitivity (Fleming, 2024). Our framework connects statistical inference in neural population codes to metacognition without making explicit assumptions about how uncertainty maps onto subjective confidence, and it quantifies a theoretical upper bound on metacognitive sensitivity attainable from population activity alone.

In human estimation tasks, the non-Gaussianity of error distributions, and in particular the long tails observed in association with higher variability, has been a source of considerable debate (Bays & Husain, 2008; Bays, 2014; Fougnie et al., 2012; Schneegans et al., 2020; Schurgin et al., 2020; Taylor & Bays, 2020; van den Berg et al., 2012; Wei & Woodford, 2025; Wilken & Ma, 2004; Zhang & Luck, 2008). Previous studies have demonstrated that models based on population coding can emulate the long tailed error distributions and also reproduce variation in subjective confidence (Bays, 2014, 2016; van den Berg et al., 2017), but the connection between the two has not previously been formalized. Here, we provide a principled account of this relationship in terms of divergence of likelihood functions which, in a translation-type model, also comprise mixture components of the estimation error distribution. We presented two complementary measures of this divergence, ℐ_loss_ and MI*ε*.

The quantity ℐ_loss_ is a measure of ancillary information that captures variability of the likelihood function in terms of Fisher information. As demonstrated by a synthetic model with Gaussian likelihoods, increasing ℐ_loss_ yields heavier tails in the marginal error distribution (see Section. 2.6). Its asymptotic limit, 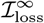 can be interpreted in terms of curvature in information geometry. Specifically, for translation-type models, Efron’s statistical curvature (Efron, 1975) is equal to the ratio between the asymptotic Fisher information loss and the Expected Fisher information (Eq. 20), establishing curvature as a formal statistical measure in a statistical submanifold.

In a recent study, Wei and Woodford (2025) emphasized the role of representational geometry in shaping error distributions, attributing deviations from Gaussianity to curvature in the neural manifold. In that framework, curvature is characterized in a more descriptive sense (Kriegeskorte & Wei, 2021), based on representational dissimilarity and low-dimensional embeddings such as principal component analysis (PCA), rather than as a formal statistical quantity. Representational dissimilarity in this framework is conceptually analogous to the “psychological scaling function” proposed to underlie error distributions in the target confusability competition (TCC) model (Schurgin et al., 2020). Note that both models incorporate Gaussian noise: the Wei and Woodford (2025) model is mathematically equivalent to the flat Gaussian noise model considered here, while the TCC model can be viewed as an approximation of it (Tomić & Bays, 2024b; Wei & Woodford, 2025).

While these models predict long-tailed behavioural error distributions, they have not connected them with information about uncertainty or addressed metacognitive sensitivity. Moreover, they assume noise-independent representational structure. In particular, the representational geometry study (Wei & Woodford, 2025) proposed that the geometry is determined by tuning functions of the neural population, largely disregarding the role of noise structure. In contrast, by formulating the statistical model as a curved exponential family, we derived analytic expressions for the curvature of the corresponding statistical submanifold and show that curvature is jointly determined by tuning and noise characteristics.

An alternative proposal for analyzing the geometry of non-Gaussian noise models is to apply an appropriate transformation to the responses that makes the variability approximately isotropic and Gaussian (Kriegeskorte & Wei, 2021; Wei & Woodford, 2025). In this study, we compared a Poisson noise model with an isotropic Gaussian noise model that approximates it via a variance-stabilizing transformation. We confirmed that the Gaussian approximation produced similar error distributions to the Poisson model, and further showed that the Expected FI was precisely equated in the two models. However, the Gaussian approximation displayed consistently lower ancillary information than the Poisson model, as measured by ℐ_loss_ or MI*e*. This suggests that the approach of transforming non-Gaussian noise models for analysis within the representational geometry framework may be sufficient for approximating first-order properties (e.g. estimation error) but could give misleading results for higher order properties (e.g. information about uncertainty). In comparison, the statistical curvature described by information geometry predicts asymptotic information loss with analytic precision, but this provides only a qualitative guide to the information loss in finite samples.

We further examined estimation and ancillary information in a Poisson population with shared fluctuations in gain, which has been proposed to describe activity in the brain more accurately (Goris et al., 2014; I.-C. Lin et al., 2015). With mean activity held constant, gain modulation had no effect on Expected FI and only a small effect on estimation error patterns, but dramatically altered ancillary information (see Section 2.7.1). Ancillary information, measured either by ℐ_loss_ or MI*e*, increased monotonically with increases in gain modulation.

We presented as an alternative measure of ancillary information the entropy-based quantity MI*e*, which characterizes variability in the likelihood in information-theoretic terms. MI*e* corresponds to the mutual information between estimation error and neural activity. Because mutual information is invariant under reparametrization, this measure provides a principled bridge between ancillary information encoded in population activity and behavioural measures of metacognitive sensitivity. From this perspective, metacognitive sensitivity in human observers can be naturally quantified by the mutual information between estimation errors *ε* and confidence reports *c*, i.e. MI(*ε*; *c*). Specifically, when confidence reports *c* are generated as a transformation of the neural activity that retains all of its ancillary information, then MI(*ε*; *c*) will equal MI*e*; otherwise MI(*ε*; *c*) will be less than MI*e*. For a given population code, MI*e* therefore quantifies an upper bound on metacognitive sensitivity attainable from the same neural representation used to generate the perceptual estimate.

Metacognitive sensitivity below this bound would imply imperfect access to uncertainty, potentially due to “metacognitive noise” (Bang et al., 2019; De Martino et al., 2013; Fleming & Daw, 2017; Guggenmos, 2022; Mamassian & de Gardelle, 2022; Maniscalco & Lau, 2014, 2016; Mueller & Weidemann, 2008; Shekhar & Rahnev, 2021; van den Berg et al., 2017), heuristic use of evidence (Maniscalco et al., 2016; Peters et al., 2017; Zylberberg et al., 2012), inference from suboptimal internal models (Herce Castañón et al., 2019), or changes in evidence distributions at perceptual level (Rahnev et al., 2012; Shekhar & Rahnev, 2024; Zylberberg et al., 2016). As we further demonstrated (see Section. 2.7.2), reduced metacognitive performance can also arise from external noise, which corrupts internal representations without contributing ancillary information. External noise could arise in sensory transduction or neural transmission upstream of the activity providing the confidence signal, or it could accumulate through recurrent connections that sustain activity over time. Note however that recurrency would also be expected to induce specific patterns of noise correlation, the consequences of which for metacognitive sensitivity remain an important subject for future research.

Alternatively, metacognitive sensitivity exceeding the bound set by MI*e* would imply access to a separate source of information about uncertainty, such as post-decisional information after a decision has been made (Fleming & Daw, 2017; Schulz et al., 2023). Dayan (2023) referred to the cases of metacognitive sensitivity being less or greater than MI*e* as *hypo-* and *hyper-*sensitivity, respectively. The quantity MI(*ε*; *c*) can be viewed as a continuous analogue of the information-theoretic measure metaℐ (Dayan, 2023), which was originally introduced to assess metacognitive sensitivity in binary discrimination tasks. Meyen et al. (2025) further interpreted meta- ℐ as ‘surplus transmitted information’, namely the additional information transmitted by a classifier’s response beyond the minimum possible transmitted information for the Type-1 performance (classification accuracy), and proposed relative metainformation (RMI) as a normalized information-theoretic measure of metacognitive sensitivity in binary tasks. Our proposed information-theoretic measure MI*e* generalizes the concept behind meta- ℐ and RMI to the domain of continuous estimation, and we have extended its application to the study of ancillary information in neural population codes.

We observed an inverted-U shape relationship between the two measures of ancillary information examined in this study (Fig. 5e). Assuming a fixed non-zero Expected FI (ℐ), MI*e* approaches zero at both extremes of ℐ_loss_ (i.e. ℐ_loss_ → 0 or ℐ_loss_ →ℐ). When ℐ_loss_ → 0, the Observed FI becomes constant across observations. In this limit, the ancillary complement represented by the Observed FI contains no variability, so conditioning on it does not reduce uncertainty in the estimation error and sensitivity thus drops to zero. Conversely, as ℐ_loss_ approaches ℐ, the likelihoods corresponding to different observations diverge strongly, with rare, very high certainty observations intermixed with frequent, very uncertain observations; the latter dominates the entropy-based measure MI*e* which falls to zero. Although all noise models exhibit similar inverted-U dependence between the two measures of ancillary information, they can differ substantially in the entropy-based ancillary information MI*e* even when matched for both Fisher information and FI-based ancillary information ℐ_loss_.

These results highlight that, for a given noise model, estimation error patterns uniquely determine the ancillary information – and thus the metacognitive sensitivity attainable for an ideal observer. However, across models, similar error patterns can arise from different noise structures that afford different amounts of ancillary information. Consequently, combining estimation errors and empirical measures of metacognitive sensitivity could help distinguish between neural models that have been considered interchangeable in previous work.

Although our framework is formulated in terms of ML estimation under an assumption of translation invariance, it admits a natural extension to Bayesian inference. For translation-type models, Fisher information is constant across parameter space, and Bayesian inference based on the uninformative Jeffreys prior (Robert et al., 2009), 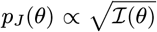, reduces to ML estimation; in this case, the Jeffreys prior is uniform. Translation invariance may be violated in more ecological settings with non-uniform stimulus statistics. In such cases, one established approach is to apply an invertible transformation of the stimulus into a sensory space where Fisher Information is uniform and the assumption of translation invariance applies (Wei & Stocker, 2015). A specific case is estimation of a scale parameter, for which the Jeffreys prior takes the form *p*_*J*_ (*σ*) ∝ 1*/σ* (Robert et al., 2009), corresponding to a uniform prior on the log scale (i.e. translation invariance in log *σ*) – a special case of the approach described by Wei and Stocker (2015) with logarithmic transformation. Extending the present framework to more general Bayesian settings that relax translational invariance and incorporate heterogeneous coding schemes will be an important direction for future work.

In summary, this work provides a principled framework relating metacognitive sensitivity to population coding of sensory evidence, offering a path toward unifying theories of perceptual processing and metacognitive computations. By formalizing how ancillary information arises from the probabilistic representation of neural codes, our approach makes concrete predictions about when and why metacognitive sensitivity diverges depending on behavioural performance, and how such divergences depend on the underlying coding characteristics. These predictions can be tested in behavioural and neurophysiological studies that concurrently collect estimation errors and confidence under controlled manipulations of set size, sensory uncertainty, prior expectations, etc. Crucially, the ability to quantify information about uncertainty directly from encoding models establishes a coherent theoretical foundation for understanding the basis of metacognition within a single formalism.

## Acknowledgements

This work is supported by International Program for Ph.D. Candidates, Sun Yat-Sen University awarded to XW, and Max Planck Society and the Humboldt Foundation awarded to PD. PMB thanks Máté Lengyel and Xue-Xin Wei for valuable discussions that inspired this work.

## Data availability statement

All code will be made publicly available upon acceptance.

## Author Contributions

**Conceptualization**: Paul M Bays

**Formal Analysis**: Xiaolu Wang, Paul M Bays

**Funding Acquisition**: Xiaolu Wang, Peter Dayan, Paul M Bays

**Investigation**: Xiaolu Wang, Paul M Bays

**Methodology**: Xiaolu Wang, Paul M Bays

**Project Administration**: Paul M Bays

**Supervision**: Paul M Bays

**Validation**: Xiaolu Wang, Peter Dayan, Paul M Bays

**Visualization**: Xiaolu Wang

**Writing – Original Draft Preparation**: Xiaolu Wang, Paul M Bays

**Writing – Review & Editing**: Xiaolu Wang, Peter Dayan, Paul M Bays

## S1 Appendix Some Mathematical Proofs

### Statistical curvature for translation-type models

We write the log-likelihood function as *l*_*θ*_ and use dot notation for derivatives with respect to *θ*, e.g. 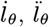 etc. For an arbitrary one-parameter statistical distribution family, we can define the covariance matrix of 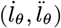, following Efron (1975, 2022), as

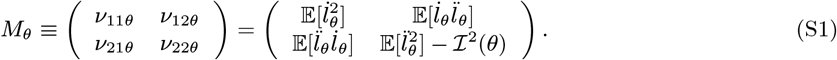

The statistical curvature at *θ* is then computed by

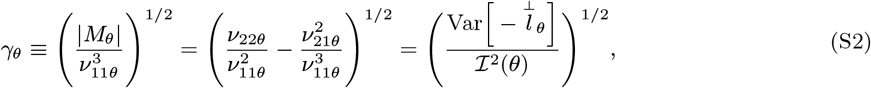

where

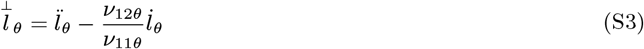

is the residual of 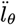 after linear regression on 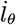. The second term vanishes for translation-type models as

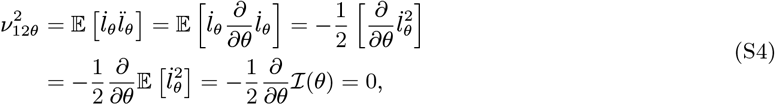

using ℐ (*θ*) = ℐ, which is invariant of *θ*. Therefore, the statistical curvature reduces to

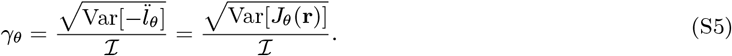

### The asymptotic loss of FI for some statistical models

For a statistical model in the form of a curved exponential family, 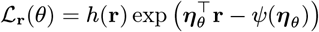, the covariance matrix of 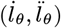 is

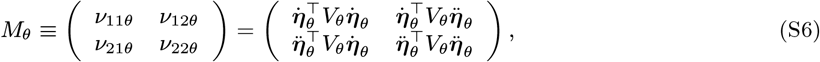

where ***η***_*θ*_ is the vector of natural parameters of the exponential family corresponding to *θ*, and *V*_*θ*_ is the covariance of **r**, which can be obtained as

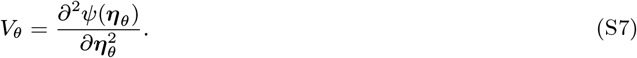

The statistical curvature defined as in Eq. S2 is in this case identical to the geometric curvature of ***η***_*θ*_ within the – generally non-Euclidean – statistical manifold of natural parameters, ℋ.

As shown in Fig. S1, exponential family models can be represented in both the sample space, ℛ, and the natural parameter space, ℋ . In (curved) exponential families, the expectation vector *µ* and natural parameter vector *η* are in a one-to-one mapping, with d*µ* = *V* d*η* (Efron, 2022). Note that both *µ* and *V* depend on *η*. For the MLE, the vector 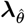 (a unit vector in the direction of 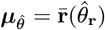 from the critical subspace *C*) is orthogonal to 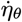 (Amari & Nagaoka, 2000). In the special case where *V* is the identity matrix, the statistical manifold in ℛ coincides exactly with that in ℛ. This is the case for the Gaussian noise model considered below.

**The Poisson model**. The likelihood function of the Poisson model can be written in the form of curved exponential families

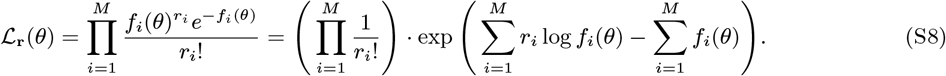

We can identify the natural parameter ***η***_*θ*_ = log **f** (*θ*) and the normalizing function *ψ*(***η***_*θ*_ ) = **f** (*θ*). The covariance of **r** is then given by

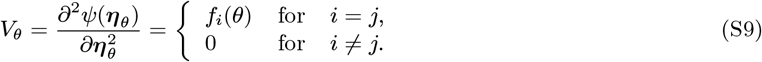

**Figure S1.**
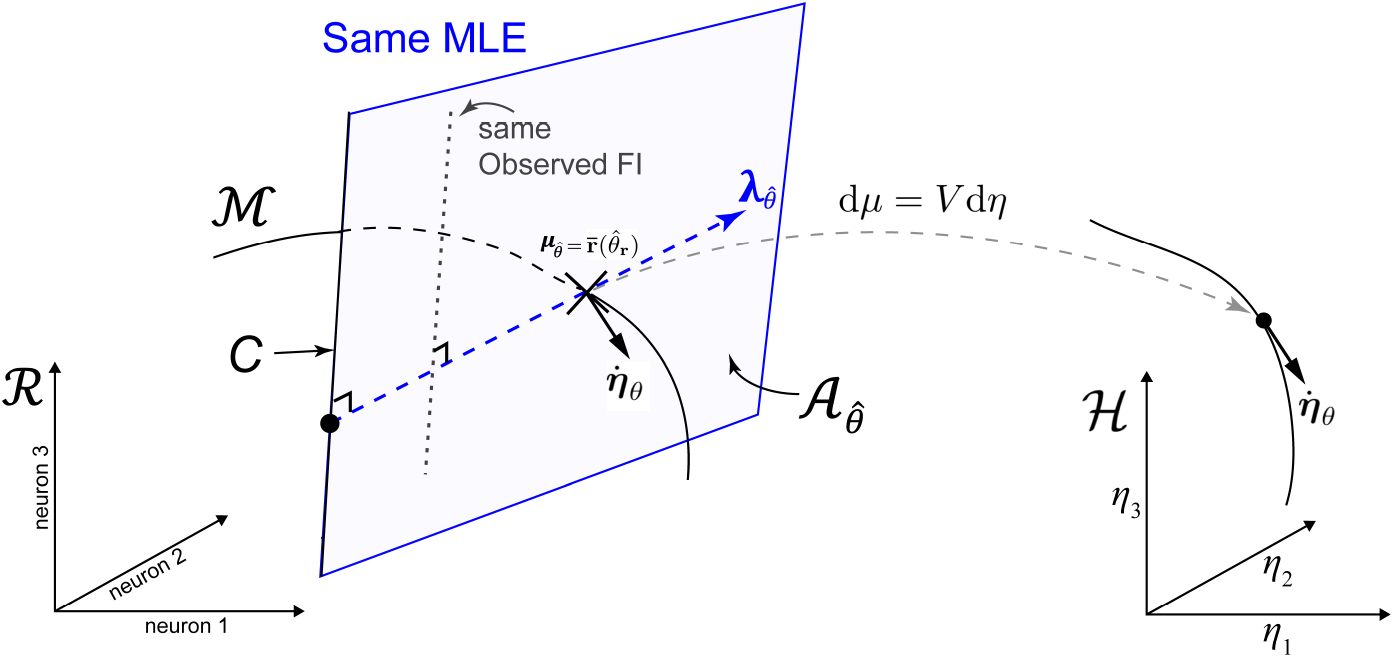
Relationship between the sample space ℛ and the natural parameter space ℋ.

Given tuning function 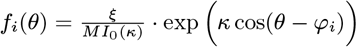, we have

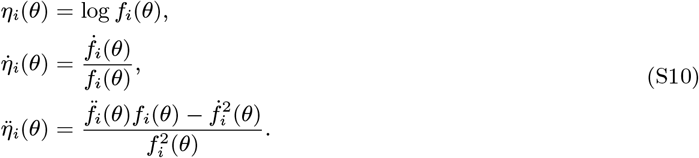

Then, the Expected FI is given by

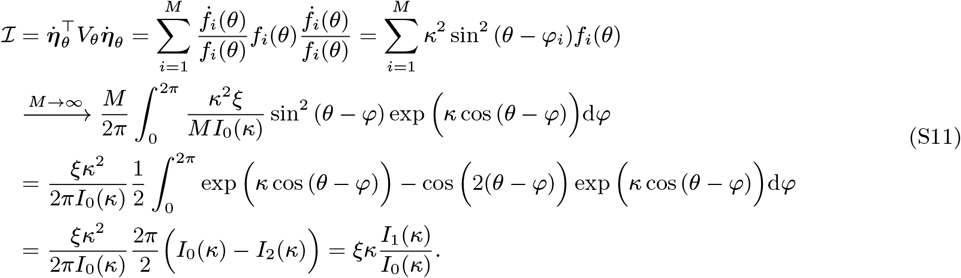

The last step uses the recurrence relations of the Bessel function of the first kind: 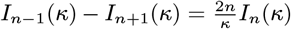.

The variance of Observed FI is given by

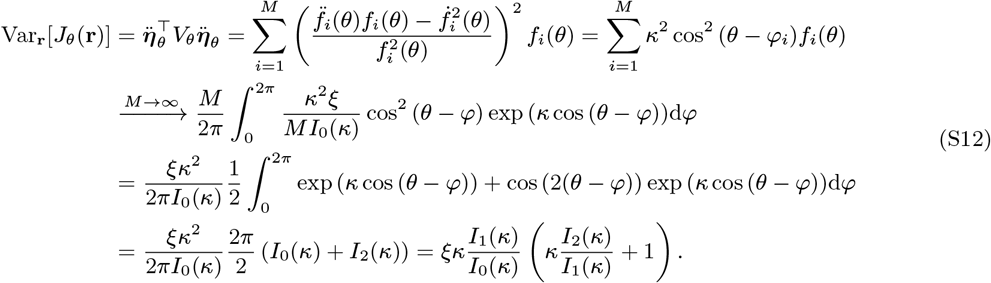

**The Gaussian model**. We can also write the likelihood function of the Gaussian model in the form of curved exponential families

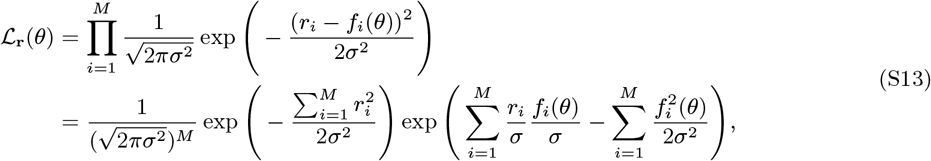

with *σ*^2^ = *ξ/M*. We can identify the natural parameter ***η***_*θ*_ = **f** (*θ*)*/σ* and the normalizing function *ψ*(***η***_*θ*_ ) = **f** (*θ*)*/*(2*σ*^2^). The covariance of **r** is then given by

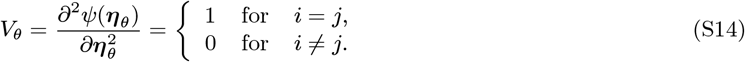

Given tuning functions 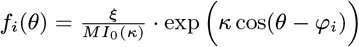, we have

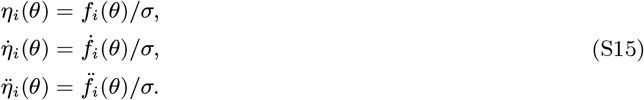

Then, we have the Expected FI

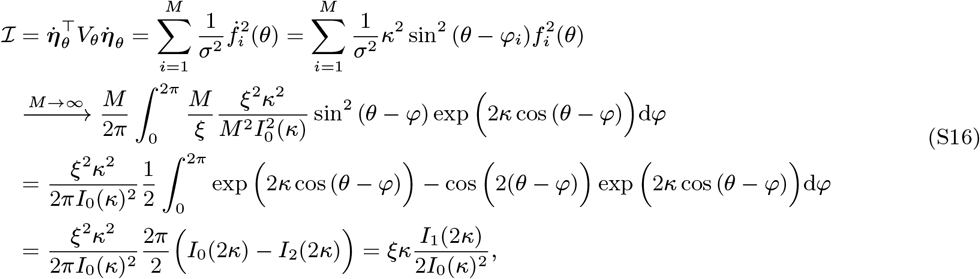

and the variance of Observed FI

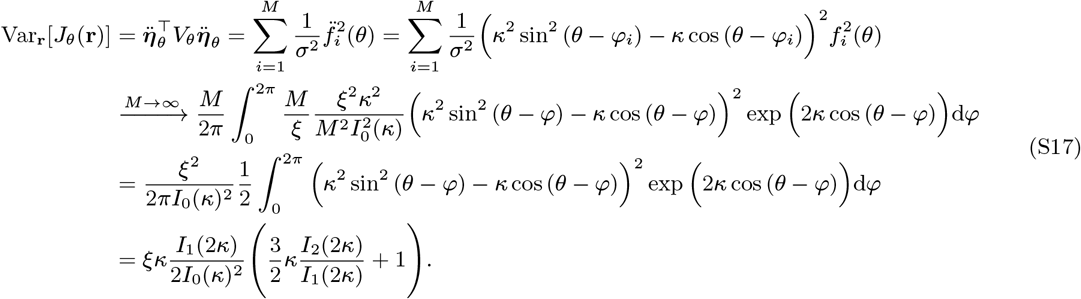

#### The modulated Poisson model

For the modulated Poisson model, the response of the *i*th neuron to stimulus *θ* is *r*_*i*_|*G* ∼ *P*(*G* · *f*_*i*_(*θ*)), where *G* ∼ Gamma(*α, s*), with 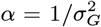 and 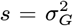. For gain *G* shared across the whole population, the likelihood function is given by

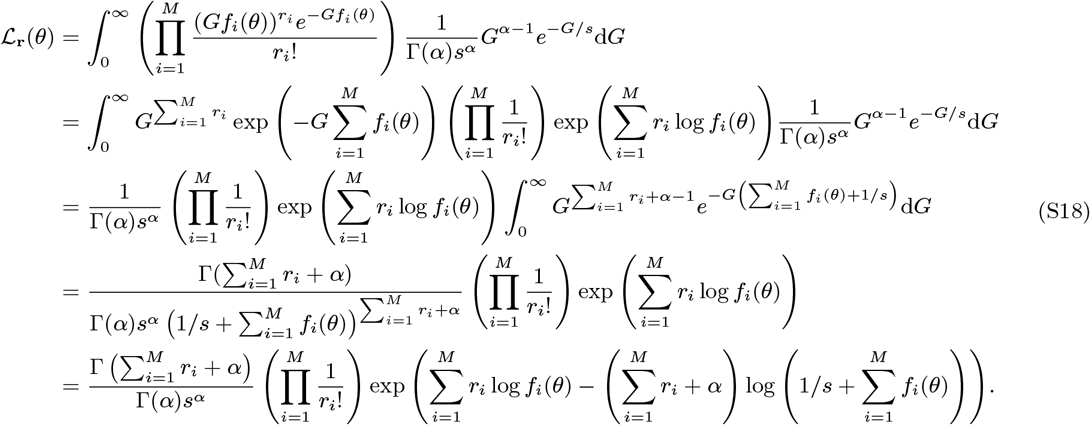

The covariance matrix of **r** is

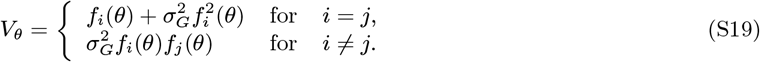

Using

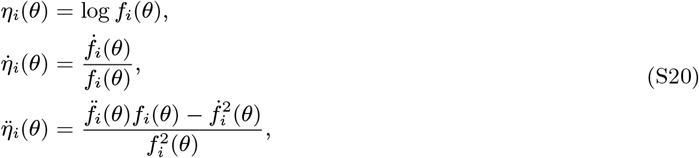

we have the Expected FI

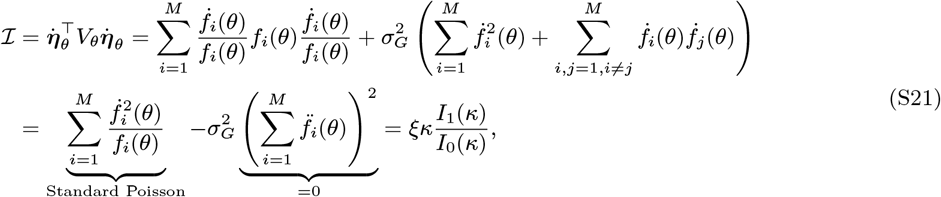

and the variance of the Observed FI

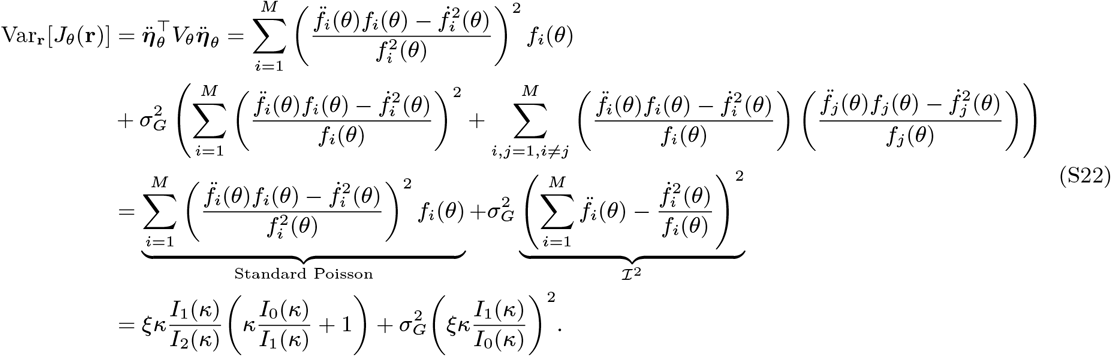

Note that for all the three noise models considered here, the Expected FI, ℐ, and the variance of the Observed FI, Var_**r**_[*J*_*θ*_(**r**)], are invariant of *θ*. Fig. S2 illustrates how the changes in ℐ, Var_**r**_[*J*_*θ*_(**r**)], and 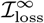 vary across different combinations of *κ* and *ξ*. When Expected FI is held constant, increasing *κ* corresponds to decreasing *ξ*.

### Matching parameters for the Poisson and Gaussian models

#### ML decoding for the Gaussian noise model

For the model with “flat” Gaussian noise, response of the *i*th of *M* neurons to a stimulus *θ* is

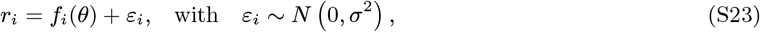

where *f*_*i*_(*θ*) is the neuron’s mean response to that stimulus.

The MLE based on responses **r** is given by

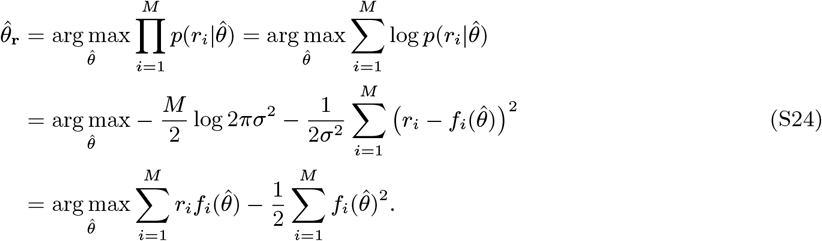

Assuming dense uniform coverage, the second term is constant and can be ignored. So we have

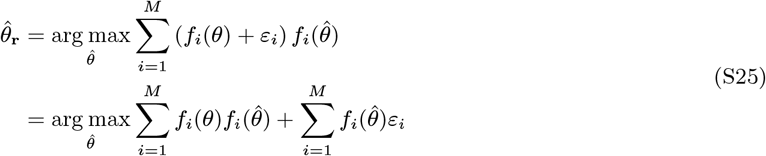

**Figure S2.**
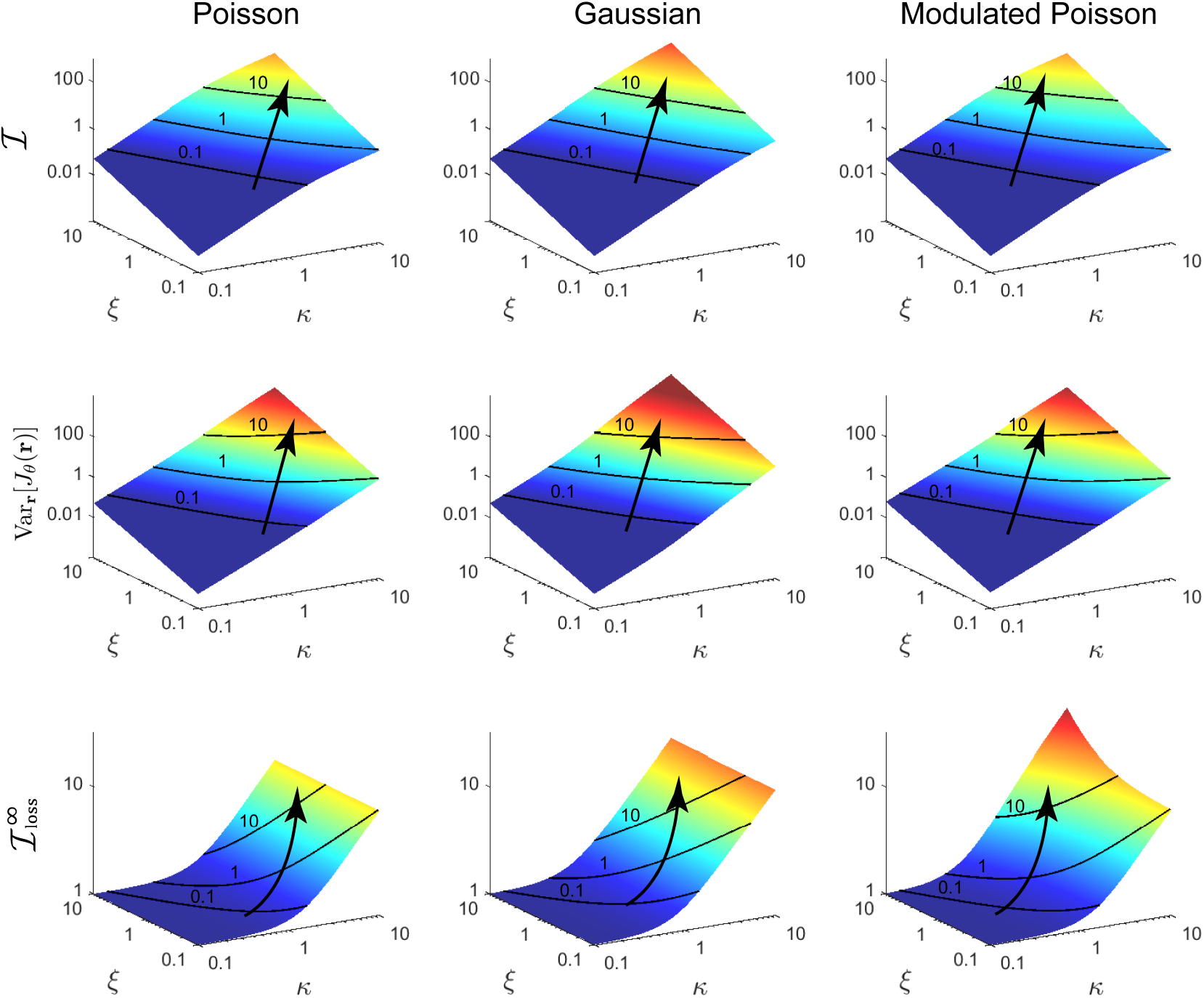
Fisher information and the asymptotic loss of FI dependent on *κ* and *ξ* for the Poisson, Gaussian, and modulated Poisson models. Contour lines represent equal Expected FI, and arrows indicate the direction Expected FI increases. The modulated Poisson is shown with 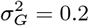.

For an infinite vector of putative estimate values 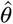, the MLE is distributed as the maximum of a multivariate normal distribution,

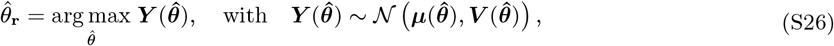

where

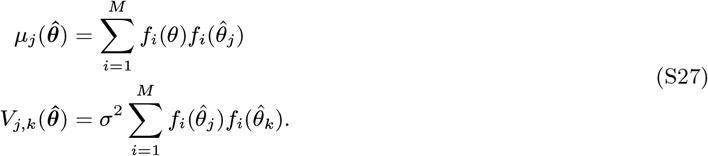

In general, this distribution cannot be obtained analytically, however it can be approximated numerically based on generating samples from the multivariate normal distribution, for which efficient methods exist.

Now let 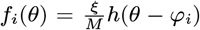 and 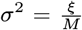, where E_*θ*_[*h*(*θ* − *φ*_*i*_)] = 1. Assuming dense uniform coverage, we have

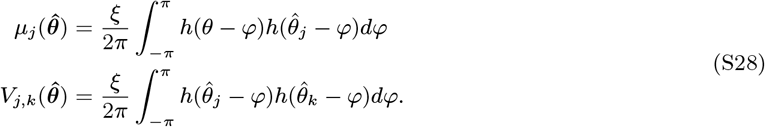

For Von Mises tuning function, *h*(*θ*) = *e*^*κ* cos(*θ*)^*/I*_0_(*κ*), we have

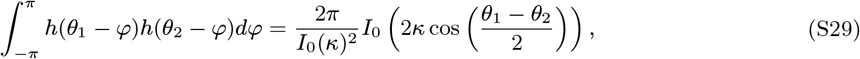

yielding,

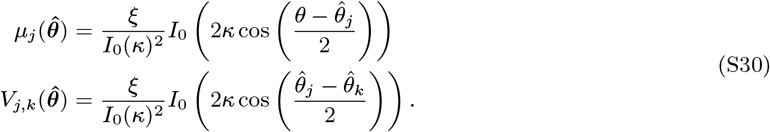

#### Variance-stabilizing transformation for the Poisson noise model

For the Poisson noise model with *r*_*i*_ | *θ* ∼ *P* (*f*_*i*_(*θ*)), we can apply a square root transformation to yield a variable 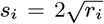 with approximately constant variance independent of *f*_*i*_(*θ*). Approximating the distribution of the transformed variable with a Gaussian gives

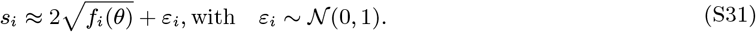

By the same argument as in the previous section, we obtain an approximate MLE of *θ* based on **s** as

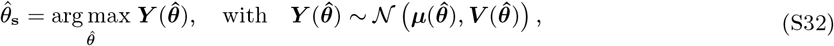

where

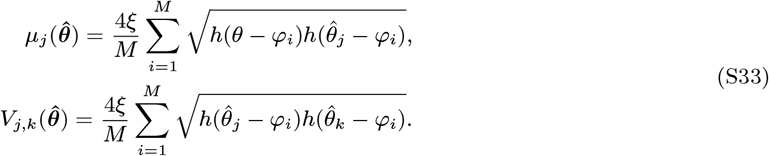

For von Mises tuning with concentration parameter *κ* and gain parameter *ξ*, and assuming dense even coverage, this yields,

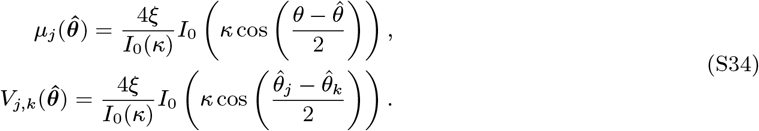

We now define

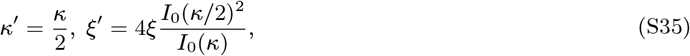

which admits the inverse mapping

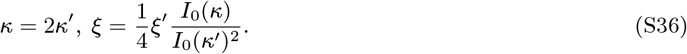

Under this reparametrization, we have

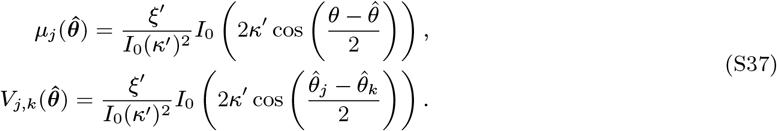

This is exactly equivalent to Eq. S30 for the Gaussian model, indicating that the Gaussian model with tuning concentration *κ*^*′*^ and gain *ξ*^*′*^ and the Poisson model with tuning concentration *κ* and gain *ξ* are approximately matched for distribution of the MLE.

